# Inhibition of carbon fixation activates ROS-linked autophagy in the single-celled green alga *Chlamydomonas reinhardtii*

**DOI:** 10.64898/2026.11.17.767326

**Authors:** Samuel Gámez-Arcas, Manuel J. Mallén-Ponce, Irene Muñoz-Palacios, Luis G Heredia-Martínez, Andrea Quintero-Moreno, Pierre Crozet, Michael Schröda, Claire Remacle, Stéphane D. Lemaire, José Luis Crespo, María Esther Pérez-Pérez

## Abstract

Autophagy is a central mechanism that promotes stress acclimation and nutrient recycling, thereby contributing to metabolic homeostasis. However, how photosynthetic carbon assimilation is integrated with autophagy remains poorly understood. Here, we establish a functional link between carbon fixation, redox signaling, and autophagy in the green microalga *Chlamydomonas reinhardtii*. Pharmacological inhibition of phosphoribulokinase (PRK), a key enzyme of the Calvin-Benson-Bassham cycle, induced chloroplast stress, reactive oxygen species (ROS) accumulation, and autophagic flux specifically in the light. Consistently, genetic loss of PRK caused light-dependent ROS accumulation and autophagy activation, which was enhanced by carbon limitation and attenuated by ROS scavenging. To determine the physiological relevance of this response, we generated an *atg8* knockout mutant and an *atg8 prk* double mutant. Loss of ATG8 increased ROS accumulation, photosynthetic dysfunction, and cell death when carbon fixation was inhibited, supporting a protective role for autophagy against light-driven metabolic stress. Although both *prk* and *atg8 prk* cells were unable to grow under high light, only *prk* cells resumed growth after transfer to permissive conditions, indicating that autophagy contributes to recovery from prolonged impairment of photosynthetic carbon metabolism. Our findings identify autophagy as a component of the regulatory network linking photosynthetic carbon assimilation with redox homeostasis and cellular acclimation to light stress.

## Introduction

Autophagy is an evolutionarily conserved catabolic process that mediates the degradation and recycling of intracellular components, thereby contributing to cellular homeostasis, nutrient remobilization, and stress acclimation in eukaryotes (He and Klionsky 2009; Mizushima et al. 2011; Liu and Bassham 2012; Marshall and Vierstra 2018). In plants and microalgae, autophagy plays a central role in coordinating cellular responses to nutrient limitation, energy deprivation and environmental stress while supporting growth, development and survival (Yoshimoto et al. 2004; Bassham et al. 2006; Perez-Perez et al. 2012b; Marshall and Vierstra 2018; Avin-Wittenberg 2019; Kajikawa et al. 2019; Signorelli et al. 2019; Mallén-Ponce et al. 2023). By recycling carbon-, nitrogen- and energy-rich cellular constituents, autophagy helps maintain metabolic homeostasis under fluctuating environmental conditions (Thompson et al. 2005; Guiboileau et al. 2013; Masclaux-Daubresse et al. 2014; Couso et al. 2018; Kajikawa et al. 2019).

Carbon metabolism is a major determinant of growth, productivity and stress acclimation in photosynthetic organisms. Through the coordinated regulation of photosynthetic carbon fixation, carbohydrate turnover, respiration and lipid metabolism, plants and microalgae continuously adjust carbon fluxes in response to environmental and developmental cues (Baena-González and Sheen 2008; Stitt and Zeeman 2012; Figueroa and Lunn 2016; Burlacot et al. 2019; Mallén-Ponce et al. 2025). Environmental stresses frequently perturb carbon allocation and energy balance, requiring extensive metabolic reprogramming to sustain growth and survival (Baena-González et al. 2007; Wang et al. 2009; Foyer et al. 2012; Lastdrager et al. 2014). Increasing evidence indicates that autophagy is intimately linked to these processes by supplying alternative respiratory substrates and facilitating metabolic flexibility during carbon or energy limitation (Perez-Perez et al. 2010; Izumi et al. 2013; Avin-Wittenberg et al. 2015; Barros et al. 2017; Couso et al. 2018; McLoughlin et al. 2020). Consistent with this view, autophagy-deficient mutants display profound alterations in amino acid, carbohydrate and lipid metabolism, as well as impaired nutrient recycling and stress tolerance (Doelling et al. 2002; Guiboileau et al. 2013; McLoughlin et al. 2018, 2020; Kajikawa et al. 2019).

Among photosynthetic model organisms, the single-celled green alga *Chlamydomonas reinhardtii* (hereafter referred to as *Chlamydomonas*) offers a versatile experimental platform for investigating the interplay between autophagy, carbon metabolism and environmental stress responses. In addition to extensive genetic and genomic resources, *Chlamydomonas* combines many of the metabolic features of higher plants with the experimental advantages of a unicellular microorganism (Merchant et al. 2007; Harris 2009; Jinkerson and Jonikas 2015; Sasso et al. 2018; Salomé and Merchant 2019). Furthermore, the conservation of most core autophagy-related proteins across the green lineage makes this microalga an excellent model for studying conserved mechanisms of autophagy regulation (Perez-Perez et al. 2010, 2012a; Kajikawa et al. 2019; Mallén-Ponce et al. 2023; Zou et al. 2025; Yang et al. 2026). A remarkable feature of *Chlamydomonas* is its metabolic flexibility, which enables it to thrive under a wide range of environmental conditions. Unlike land plants, which rely predominantly on photoautotrophic growth, *Chlamydomonas* can grow photoautotrophically using CO₂ as its sole carbon source, mixotrophically in the presence of both light and reduced carbon sources (acetate or glycolate), or heterotrophically in the dark when acetate is supplied as an external carbon source (Harris 2009; Johnson and Alric 2013; Sasso et al. 2018; Salomé and Merchant 2019; Cavallari et al. 2026). The capacity to utilize acetate provides a unique opportunity to manipulate carbon availability independently of photosynthetic carbon fixation, as photosynthetic machinery mutants are viable, making *Chlamydomonas* a versatile model for studying carbon allocation. This metabolic versatility has also enabled the study of nutrient-sensing pathways, such as TOR signaling, and their interplay with autophagy and carbon status (Perez-Perez et al. 2010; Mallén-Ponce et al. 2022, 2025).

In both plants and microalgae, nutrient deprivation frequently triggers metabolic reprogramming, including alterations in carbon partitioning, accumulation of storage compounds and activation of autophagy. At the same time, perturbations in carbon metabolism often result in increased production of reactive oxygen species (ROS), particularly in chloroplasts and mitochondria (Apel and Hirt 2004; Foyer and Noctor 2009; Mittler 2017). Although ROS act as essential signaling molecules, their excessive accumulation can damage proteins, lipids and nucleic acids, making redox homeostasis a critical determinant of cellular fitness (Noctor et al. 2018; Waszczak et al. 2018; Wakao and Niyogi 2021).

Autophagy has emerged as a crucial component of cellular antioxidant defense and quality-control systems. Oxidative stress induces autophagy, whereas impaired autophagic activity frequently leads to elevated ROS levels and enhanced sensitivity to environmental stress (Xiong et al. 2007; Yoshimoto et al. 2009; Perez-Perez et al. 2012a; Signorelli et al. 2019). Beyond the degradation of oxidized proteins, autophagy contributes to the selective removal of damaged chloroplasts and other organelles, thereby limiting ROS production and preserving organellar function (Woodson 2019; Otegui et al. 2024). Besides its role in the removal of irreversibly damaged cellular components, autophagy is itself subject to tight redox regulation. Accumulating evidence indicates that several components of the core autophagy machinery, including ATG4 and ATG3, act as redox-sensitive proteins, enabling ROS signals to directly modulate autophagic activity (Scherz-Shouval et al. 2007; Perez-Perez et al. 2014, 2016; Woo et al. 2014; Frudd et al. 2018; Mallén-Ponce and Pérez-Pérez 2023).

Current evidence supports the view that autophagy, carbon metabolism and oxidative stress constitute an integrated regulatory network that governs cellular acclimation to changing environmental conditions. Beyond its established role in nutrient recycling, autophagy contributes directly to cellular carbon homeostasis by sustaining photosynthetic performance, promoting metabolic remodeling and supporting the recycling of cellular components for efficient carbon assimilation. Accordingly, autophagy-deficient mutants frequently display altered carbon fluxes, impaired starch and amino acid metabolism, reduced respiratory flexibility and decreased fitness under carbon-limiting conditions (Guiboileau et al. 2013; Masclaux-Daubresse et al. 2014; Avin-Wittenberg et al. 2015; Barros et al. 2017; McLoughlin et al. 2020). Moreover, selective autophagy contributes to chloroplast quality control, thereby preserving photosynthetic competence and maintaining carbon fixation capacity (Izumi et al. 2017; Otegui 2018; Woodson 2019). Therefore, autophagy should be viewed not only as a recycling pathway activated during nutrient or carbon limitation, but also as an active regulator of carbon-use efficiency, metabolic allocation and physiological acclimation in photosynthetic organisms. In this context, the metabolic flexibility of *Chlamydomonas*, including its ability to utilize acetate as an alternative carbon source, provides an effective experimental framework to dissect the molecular connections between carbon assimilation, nutrient signaling, redox regulation and autophagy.

In this study, we investigate the links between carbon assimilation, autophagy and oxidative stress in *Chlamydomonas*. Using both genetic and pharmacological approaches to impair phosphoribulokinase (PRK) activity, we show that reduced Calvin-Benson-Bassham (CBB) cycle function triggers autophagy-related responses and profound physiological adjustments. Our findings support the notion that autophagy is not only a consequence of metabolic stress but also an important component of the cellular acclimation mechanisms that help maintain carbon-use efficiency and redox homeostasis in photosynthetic organisms.

## Materials and Methods

### Strains, media, and growth conditions

*Chlamydomonas reinhardtii* WT strain 4A+ (WT4A+; CC4051), the Clip Library WT strain (CC5325) and the *prk* mutant ΔPRK (LMJ.RY0402.119555) were obtained from the *Chlamydomonas* Resource Center (http://www.chlamycollection.org). The *atg8* mutant and the complemented strain expressing StrepII-tagged ATG8 (*atg8*::pStrep-ATG8) were generated in this study as described below. Other *Chlamydomonas* strains used in this work are listed in Supplemental Table S1. *Chlamydomonas* cells were grown under low light (LL; 20 µmol photons m^-2^ s^-1^), medium light (ML; 50 µmol photons m^-2^ s^-1^), high light (HL; 500 µmol photons m^-2^ s^-1^) or in darkness on an orbital shaker (100 rpm) at 25°C in Tris-acetate-phosphate (TAP) or Tris-phosphate (TP) medium, as previously described (Harris, 1989). When required, cells in the exponential growth phase (∼10^6^ cells ml^-1^) or stationary growth phase (∼10^7^ cells ml^-1^) were treated with glycolaldehyde (GLA; Sigma-Aldrich, G6805) or N-acetyl-cysteine (NAC; Sigma-Aldrich, A7250) at the indicated concentrations for the specified times. Fresh acetate (Panreac, 141008.1212) was added to the *Chlamydomonas* culture when indicated. For growth on solid medium, 1.2% Bacto agar (Difco, 214010) was added to the culture medium.

### Protein preparation from *Chlamydomonas* cells

*Chlamydomonas* cells from liquid cultures were collected by centrifugation at room temperature (4000 x *g,* 4 min), washed with lysis buffer 1 (50 mM Tris-HCl (pH 7.5)), and resuspended in a minimal volume of the same buffer. Cells were lysed by two cycles of slow freezing at -80°C followed by thawing at room temperature. The soluble protein extract was separated from the insoluble fraction by centrifugation at 15000 x *g* for 20 min at 4°C in a microcentrifuge. For the analysis of membrane-associated proteins, such as VIPP1 and VIPP2, cells were collected by centrifugation, washed with lysis buffer 1, frozen in liquid nitrogen, and stored at - 80°C until use. Cell pellets were lysed by incubation at 37°C for 30 min in lysis buffer 2 (50 mM Tris-HCl (pH 7.5), 2% (w/v) SDS, 10 mM EDTA, 1x protease inhibitor cocktail from Roche, 4693116001). The total protein extract was obtained by centrifugation at 15000 x *g* for 20 min at 4°C in a microcentrifuge. Protein concentration was determined using either the Coomassie dye-binding assay (BioRad, 500-0006) or the bicinchoninic acid (BCA) protein assay (Supelco, B9643), according to the manufacturerś instructions.

### Immunoblot analysis

For immunoblot analyses, total protein extracts (5 to 20 μg) were separated by SDS-PAGE using 12% (v/v) gels for RPS6 detection, or 15% (v/v) gels for FKBP12, RPL37, VIPP1, VIPP2, PRK and ATG8 detection, and subsequently transferred to 0.45 μm nitrocellulose membranes (Cytiva; 10600003). Primary antibodies raised against ATG8 (Perez-Perez et al. 2010), FKBP12 (Crespo et al. 2005), RPS6 (Couso et al. 2020), RPL37 (Agrisera; AS122115), VIPP1 (Liu et al. 2005) and VIPP2 (Theis et al. 2019), PRK (Boisset et al. 2023) were diluted 1:3.000, 1:5.000, 1:2.000, 1:10.000, 1:2.000, 1:2.000 and 1:10.000, respectively, in phosphate-buffered saline (PBS) containing 0.1% (v/v) Tween-20 (Applichem, A4974) and 5% (w/v) nonfat dry milk (Applichem, A0830). Horseradish peroxidase-conjugated anti-rabbit secondary antibody (Sigma-Aldrich, A6154) was diluted 1:10.000 in the same solution. Proteins were detected using the Luminata Crescendo Western HRP substrate (Millipore, WBLUR0500) and visualized with an ImageQuant 800 imaging system (Amersham). When required, protein bands were quantified using the ImageQuant TL software (Amersham).

### Chlorophyll quantification

Chlorophyll pigments were extracted from *Chlamydomonas* cells grown under the conditions described above and quantified spectrophotometrically as described by (Porra et al. 1989). Chlorophyll *a* (Chl *a*) and chlorophyll *b* (Chl *b*) concentrations (μg/ml) were calculated using the equations reported by (Porra et al. 1989) based on absorbance measurements at 663 and 646 nm, with absorbance at 750 nm used as a baseline reference to correct for turbidity and light scattering.

### Chlorophyll fluorescence measurement

Photosynthetic parameters were measured using a Dual-PAM-100 fluorometer (Walz). *Chlamydomonas* cells grown autotrophically in TP medium under the indicated conditions were treated with GLA for the specified times. Subsequently, 2 ml of culture (∼10^6^ cells ml^−1^) were used to determine the maximum quantum efficiency of PSII photochemistry (*F*_v_/*F*_m_) and the relative electron transport rate (rETR). For *F*_v_/*F*_m_ measurements, cells were dark-adapted for 10 min under continuous stirring to determine the minimum fluorescence yield (*F*_o_). Maximum fluorescence yield (*F*_m_) was then measured in the presence of 20 μM DCMU under red actinic light (25 µmol photons m^-2^ s^-1^). Variable fluorescence (*F*_v_) was calculated as *F*_m_ – *F*_o_, and the maximum photochemical efficiency of PSII was expressed as the *F*_v_/*F*_m_ ratio. To determine the effective quantum yield of PSII (Y(II)) and rETR, cells were dark-adapted for 10 min under continuous stirring and then exposed to light (50 µmol photons m^-2^ s^-1^) for 5 min. Y(II) at 50 μmol photons m^-2^ s^-1^ and rETR values were calculated using the Dual-PAM software according to manufactureŕs instructions. NADPH fluorescence was simultaneously monitored using the NADPH module integrated into the Dual-PAM-100 system (Walz). Cells were dark-adapted for 10 min before measurement, and fluorescence changes were recorded during dark-to-light transitions under the indicated illumination conditions. NADPH fluorescence traces were analyzed using Dual-PAM software according to the manufactureŕs recommendations.

### Oxygen evolution and consumption measurements

Oxygen evolution and consumption rates were determined at 25°C using a Clark-type oxygen electrode (Chlorolab 2+ System; Hansatech, Norfolk,UK). *Chlamydomonas* WT, *prk* and *atg8* cells grown under the indicated conditions were harvested by centrifugation, resuspended in fresh medium, and adjusted to the desired cell density. Measurements were performed in a thermostated reaction chamber containing 1 ml of cell suspension under continuous stirring. Prior to measurements, cells were dark-adapted for 5 min. Oxygen exchange was then monitored during illumination with white light (50 µmol photons m^-2^ s^-1^) for 5 min, followed by a 5 min dark period. Photosynthetic activity was calculated as the difference between the rate of oxygen evolution in the light and the rate of oxygen consumption in the dark.

Oxygen exchange rates were normalized to culture density measured as absorbance at 750 nm (OD_750nm_) and expressed as µmol O_2_ min^-1^/OD_750nm._

### Viability assay

Cell viability was assessed by determining the percentage of *Chlamydomonas* WT and *atg8* cells grown under the indicated conditions, as previously described in (Perez-Martin et al. 2014). Briefly, cells were stained with Evans blue dye (E2129; Sigma), which selectively stains nonviable cells. A 450 μl aliquot of *Chlamydomonas* culture was incubated with 0.1% (w/v) Evans blue for 5 min, washed with 500 μl of TAP medium, and resuspended in an equal volume of TAP medium. Cells were then examined using phase-contrast microscopy to visualize dye uptake, with Evans blue-positive cells scored as nonviable.

### ROS determination

Intracellular ROS levels were determined in *Chlamydomonas* WT, *prk* and *atg8* cells grown under the indicated conditions using a Cary Eclipse fluorescence spectrophotometer (Agilent Technologies). ROS measurements were performed as previously described by (Perez-Perez et al. 2012a), following the method of (Joo et al. 2005). Briefly, total protein extracts were prepared as described above, and aliquots of the corresponding cell cultures were incubated with 2′,7′-dichlorodihydrofluorescein diacetate (H_2_DCFDA; Invitrogen, D-399). Upon oxidation by intracellular ROS, H_2_DCFDA is converted into the fluorescent compound 2’,7’-dichlorofluorescein (DCFDA), whose fluorescence intensity was used as an indicator of ROS accumulation. Relative fluorescence units were normalized to the reference sample indicated in each experiment.

### roGFP fluorescence recordings

roGFP fluorescence was recorded using a Cary Eclipse fluorescence spectrophotometer as previously described for *Chlamydomonas* by (Niemeyer et al. 2021). The *Chlamydomonas* strains expressing the H_2_O_2_ sensor roGFP2-Tsa2ΔC_R_ targeted to either the cytosol or the chloroplast are listed in Supplemental Table S1. Prior to fluorescence measurements, cultures were treated with 1 mM GLA for the indicated time periods. Fluorescence emission was monitored at 510 nm following sequential excitation at 405 and 488 nm, and the redox state of the sensor was expressed as the fluorescence ratio (405/488). The untransformed parental strain UVM4, lacking roGFP expression, was processed under identical conditions and used as a negative control to define baseline cellular autofluorescence (set as zero). To evaluate probe oxidation and relative subcellular ROS generation, autofluorescence values obtained from UVM4 cells were subtracted from the corresponding fluorescence signals of the cytosolic- and chloroplast-targeted lines. Data are expressed as arbitrary units (A.U.) relative to the UVM4 baseline, representing relative changes in probe oxidation across subcellular compartments.

### Metabolomic analysis

Metabolite extraction and analysis were performed as previously described (Mallén-Ponce et al. 2025). Briefly, *Chlamydomonas* cells grown under the indicated conditions were harvested by centrifugation (4000 x *g*, 2 min, 4°C). Cell pellets were immediately frozen in liquid nitrogen and stored at -80°C until further processing. Frozen pellets were resuspended in ice-cold methanol:chloroform (70:30; v/v) and homogenized by 15 cycles of 30 s vortexing followed by immersion in liquid nitrogen to maintain samples at low temperature throughout the extraction procedure. Subsequently, Milli-Q water was added to induce phase separation, and samples were centrifugated (15 000 x *g*, 20 min, 4°C). The upper polar phase was transferred to a fresh tube and dried overnight in a vacuum concentrator. Dried extracts were resuspended in 50 μl of Milli-Q water and analyzed by LC-MS/MS using a QTRAP 6500+ mass spectrometer (Sciex) at the Chromatography Facility of the Instituto de Bioquímica Vegetal y Fotosíntesis (IBVF). Metabolites were identified and quantified based on chromatographic and mass spectrometric parameters as previously described (Mallén-Ponce et al., 2025). Peak areas were normalized to the internal standard acetaminophen.

### atg8 mutant generation by CRISPR/Cas9 technology

The *ATG8* gene (accession number: Cre16.g689650) sequence was retrieved from the Phytozome database (https://phytozome-next.jgi.doe.gov). A sgRNA targeting *ATG8* (5’-CCGACCCCCGACATTCAAGCAGG-3’) and the Cas9 protein were designed as previously described (Shin et al. 2016) and obtained from LABOMICS (Belgium). The *Chlamydomonas* WT4A+ strain was co-transformed by electroporation with the *ATG8* sgRNA/Cas9 ribonucleoprotein (RNP) complex and the pSL18 plasmid, carrying the *APHVIII* gene that confers resistance to paromomycin. Briefly, exponentially growing cells in TAP medium (approximately 1 x 10^6^ cells ml^-1^) were harvested and concentrated 100-fold in TAP supplemented with 60 mM sucrose. Subsequently, 250 μl of concentrated cells were mixed with 1 μg of linearized pSL18 plasmid and incubated on ice for 15 min. Next, 25 μg sgRNA and 20 μg Cas9 were added to the cell suspension and incubated for an additional 5 min on ice. For RNP assembly, prior to the incubation with cells and pSL18 mixture, sgRNA and Cas9 protein were incubated together for 10 min at 25°C followed by 30 min at 37°C. The mixture containing cells, pSL18 linearized plasmid, sgRNA and Cas9 was transferred to a 0.4 cm gap electroporation cuvette (BioRad, 1652081) and subjected to electroporation using a Gene Pulser Xcell system (BioRad) under the following conditions: 800 V (2000 V/cm), 25 μF capacitance, and no shunt resistance. Immediately after electroporation, cells were allowed to recover in TAP medium supplemented with 60 mM sucrose for 16 h under low light (20 μmol photons m^-2^ s^-1^) with gentle agitation (50 rpm). Cells were then plated on TAP-agar plates supplemented with 100 μg/μl paromomycin. Resistant colonies were subjected to successive rounds of selection using increasing concentrations of paromomycin. Finally, the selected colonies were analyzed by immunoblot using a CrATG8 specific antibody as described above. Those colonies lacking detectable ATG8 protein accumulation were chosen for further molecular and genetic characterization.

### *atg8* genomic analysis

Genotyping of the *ATG8* locus was performed by PCR using primer pairs specific for the *ATG8* allele or combination of *ATG8* and *APHVIII* (paromomycin-resistance cassette) specific primers to detect the disrupted *ATG8* allele. The *ATG8* allele was amplified using ATG8 specific forward and reverse primers, whereas the mutant *ATG8* allele was scored using a combination of *ATG8* and paromomycin-resistance primers (Supplemental Table S2). PCR reactions were carried out in a final volume of 50 µl containing Taq DNA polymerase in the presence of 1X Taq DNA polymerase buffer, 1 µM of each primer, 0.5 mM dNTPs, 1 µl genomic DNA, and 2.5 µl DMSO, according to the manufactureŕs recommendations. The PCR was performed under the following conditions: an initial denaturation step at 95°C for 2 min, followed by 35 cycles of 95°C for 1 min, 60°C for 1 min, and 72°C for 1 min, with a final extension step at 72°C for 10 min.

### RNA isolation and RT-qPCR analysis

*Chlamydomonas* WT and *atg8* cells grown in liquid cultures were harvested by centrifugation (4000 x *g*, 4 min) and washed once with 50 mM Tris-HCl (pH 7.5). Cell pellets were immediately frozen in liquid nitrogen and stored at -80°C until use. Total RNA was isolated from frozen cell pellets as previously described (Crespo et al. 2005). First-strand cDNA was synthesized from 1 μg of total RNA using the iScript cDNA Synthesis Kit (BioRad, 170-8891) according to manufactureŕs instructions, in a final reaction volume of 20 μl. Quantitative real-time PCR (RT-qPCR) was performed using an iCycler iQ Real-Time PCR Detection System (Bio-Rad). Amplification reactions were carried out in a final volume of 18 μl using the SsoAdvanced Universal SYBR Green Supermix (BioRad; 172-5271) and gene specific primers (Supplemental Table S2). The gene expression data were normalized to *CBLP* (Cre06.g278222) expression, a constitutively expressed gene encoding a protein homologous to the β-subunit of a G-protein, serving as an internal reference gene used as control.

### Complementation of *atg8* mutant using StrepII-tagged *ATG8*

The *Chlamydomonas ATG8* coding sequence was synthesized by GeneCust Europe (http://www.genecust.com) and cloned into the pAGM8035 recipient vector using the Modular Cloning (MoClo) strategy as previously described (Crozet et al. 2018). The resulting plasmid, named pM65, contained the *ATG8* coding sequence fused to an N-terminal StrepII tag and placed under the control of the *PSAD* promoter and terminator. In addition, the construct carried a hygromycin-resistance cassette for selection in *Chlamydomonas*. The recipient plasmid pAGM80359, promoter, terminator, StrepII tag and antibiotic-resistance modules were obtained from the Chlamy-MoClo toolkit (Addgene) as described by Crozet et al. (2018). All restriction and ligation reactions were performed using *Bbs*I-HF or *Bsa*I-HF restriction enzymes (New England Biolabs) and T4 DNA ligase (New England Biolabs) in a CutSmart buffer supplemented with 1 mM ATP, following standard MoClo procedures. After verification of DNA integrity, 500 ng of pM65 plasmid were introduce into the *atg8* mutant by electroporation as described above. Following transformation, cells were plated on TAP-agar medium supplemented with hygromycin B (20 μg/μl). Resistant colonies were subjected to successive rounds of selection on increasing hygromycin concentrations. Transformants were screened by immunoblot analysis using anti-CrATG8 antibodies for confirmation. A complemented line showing stable expression of StrepII-ATG8 was selected for further analysis and designated *atg8*::Strep-ATG8.

### *atg8* prk double mutant generation by genetic crosses

Genetic crosses of single *atg8* and *prk* mutants were performed as described in (Duby and Matagne 1999). In summary, zygotes were matured for 3-4 days under continuous light on nitrogen-free TP agar plates. After maturation, blocks of agar carrying 50-100 zygotes were transferred to fresh TAP agar plates and exposed to chloroform vapor for 30 s. Germination was induced by exposure to light and restoration of nitrogen. After 10 days, 150-300 meiotic clones were sampled and further analyzed by antibiotic selection and immunoblot tests.

### Statistical Analysis

Data were obtained from at least three independent biological replicates and analyzed using GraphPad Prism software. Statistical significance was assessed using two-tailed Student’s *t*-tests for pairwise comparisons and one-way analysis of variance (ANOVA) for comparisons involving more than two groups. For all representations, data are presented as mean ± SD unless otherwise stated. Differences were considered statistically significant at *P*<0.05. Statistical significance is indicated as follows: *P*<0.05 (*), *P*<0.01 (**), and *P*<0.001 (***). A detailed summary of the statistical analyses performed for experiments shown in the main and supplementary figures is provided in Supplemental Table S3 and Supplemental Table S4, respectively.

## Results

### Inhibition of carbon assimilation activates autophagy

Given the fundamental role of autophagy in maintaining cellular homeostasis under nutrient limitation, we investigated whether carbon assimilation regulates this catabolic process in *Chlamydomonas*. To this aim, we monitored autophagy in wild-type (WT) *Chlamydomonas* cells treated with glycolaldehyde (GLA), a specific PRK inhibitor that blocks CO_2_ fixation and the CBB cycle (Fig. 1A) (Takahashi and Murata 2005; Mallen-Ponce et al. 2022). First, we tested the sensitivity of WT cells to the presence of 0.5 mM GLA under different trophic regimes, using CO_2_ (TP medium) or both CO_2_ and acetate (TAP medium) as carbon sources under moderate light (50μmol photons m^-2^ s^-1^). We observed that GLA impaired cell growth in autotrophic conditions, but this effect was largely attenuated in the presence of acetate (Supplemental Fig. S1A). To further confirm the negative effect of GLA on carbon fixation, we analyzed the abundance of different metabolites from the CBB cycle in GLA-treated cells. As expected, the level of the PRK substrate ribulose-5-phosphate (Ru5P) and its interconvertible isomer xylulose-5-phosphate (Xu5P) strongly increased within 30 min of GLA addition and stayed upregulated along treatment (Supplemental Fig. S1B), indicating that PRK activity was efficiently blocked.

**Figure 1.**
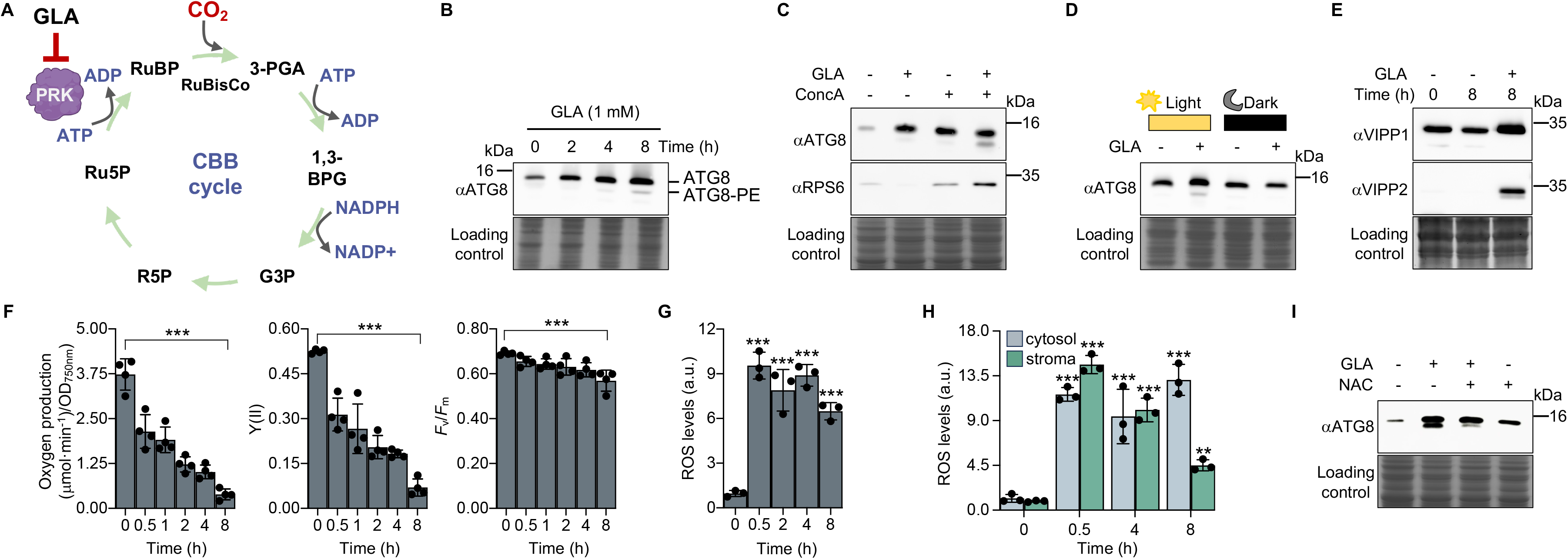
Inhibition of the Calvin Benson Bassham (CBB) cycle activates autophagy in *Chlamydomonas*. **(A)** Schematic representation of CO_2_ fixation, indicating the reaction step blocked by glycolaldehyde (GLA). GLA inhibits PRK, a key enzyme of the CBB cycle that catalyzes the conversion of ribulose-5-phosphate to ribulose-1,5-bisphosphate, thereby affecting CO_2_ assimilation and downstream carbon metabolism. Ru5P, ribulose-5-phosphate; RuBP, ribulose-1,5-bisphosphate; 3PGA, 3-phosphoglycerate; 1,3-BPG, 1, 3 bisphosphoglycerate; G3P, glyceraldehyde-3-phosphate; R5P, ribose-5-phosphate. **(B)** Immunoblot analysis of ATG8 protein levels in WT *Chlamydomonas* cells grown autotrophically under medium light (50 μmol photons m^-2^ s^-1^) conditions and treated with 1 mM GLA for 0, 2, 4 and 8 h. **(C)** Autophagic flux assay in GLA-treated cells. Immunoblot analysis of ATG8 protein (upper panel) and the ribosomal protein RPS6 (middle panel) in WT cells treated (+) with 1 mM GLA and/or 0.1 μM concanamycin A (ConcA) for 8 h. **(D)** Immunoblot analysis of ATG8 protein levels in WT cells treated with 1 mM GLA for 8 h under light or dark conditions. **(E)** Immunoblot analysis of VIPP1 (upper panel) and VIPP2 (middle panel) proteins in WT cells treated with 1 mM GLA for 8 h. **(F)** Photosynthetic parameters, including oxygen evolution, effective quantum yield of PSII (Y(II)), and maximum quantum efficiency of PSII (*F*_v_/*F*_m_), measured in WT cells before (0 h) and after treatment with 1 mM GLA for 0.5, 1, 2, 4 and 8 h. **(G)** ROS levels in WT cells treated with 1 mM GLA for 0, 0.5, 2, 4 and 8 h. **(H)** ROS levels in different subcellular compartments (cytosol and stroma) of WT cells treated with 1 mM GLA for 0, 0.5, 4 and 8 h. **(I)** Immunoblot analysis of ATG8 protein levels in WT cells treated with 1 mM GLA and/or 10 mM N-Acetyl-L-Cysteine (NAC) for 8 h under light conditions. Experiments shown in (*B*, *C*, *E*-*I*) were performed under standard growth light conditions (50 μmol photons m^-2^ s^-1^), whereas (*D*) was conducted either in the light or in the dark. Coomassie Brilliant Blue stained gels (lower panels) were used as loading controls in (*B-E, I*). Error bars represent SD of the mean values. In (*G*) and (*H*), ROS levels are expressed in arbitrary units (a.u.), with the untreated sample used as reference. Asterisks indicate statistically significant differences determined by one-way ANOVA followed by Bonferronís test: \*\*\**P*<0.001; \*\**P*<0.01; \**P*<0.05; and n.s. not significant ≥0.05. Data represent the mean of at least three independent biological experiments. Details of the statistical analyses are provided in Supplemental Table S3.

Next, we analyzed ATG8 lipidation, a hallmark of autophagy activation (Perez-Perez et al. 2010; Klionsky et al. 2021), in GLA-treated *Chlamydomonas* cells under autotrophic growth. GLA treatment resulted in an increase in both free and lipidated ATG8 (ATG8-PE) (Fig. 1B). We found that 1 mM GLA was sufficient to activate autophagy (Supplemental Fig. S2A). The effect of GLA on the abundance of ATG8 was observed within 2-4 h of drug treatment, being ATG8-PE clearly detected in 4-8 h (Fig. 1B). We also determined autophagic flux in GLA-treated cells using concanamycin A (ConcA), an inhibitor of vacuolar ATPase (Drose et al. 1993). We have previously shown that ConcA blocks autophagic flux in *Chlamydomonas*, which prevents the degradation of the ribosomal proteins RPS6 and RPL37 and triggers the accumulation of ATG8 and ATG8-PE (Couso et al. 2018). Our results showed that GLA activates autophagic flux in *Chlamydomonas*. GLA treatment downregulated the abundance of RPS6 while the presence of ConcA blocked the degradation of this protein (Fig. 1C). Remarkably, we found that GLA triggered autophagy in the light but not in the dark (Fig. 1D), suggesting that the activation of this degradative process by GLA is linked to the inhibition of photosynthetic assimilation of CO_2_. We also investigated whether GLA treatment may lead to chloroplast stress. For this, we analyzed the level of VIPP (Vesicle-inducing protein in plastids) proteins. VIPP1 and VIPP2 aid thylakoid biogenesis and membrane maintenance in microalgae (Nordhues et al. 2012), and both proteins strongly accumulate under various chloroplast-related stress conditions including high light (Nordhues et al. 2012), ClpP protease depletion (Ramundo et al. 2014) and inhibition of fatty acid synthesis (Heredia-Martinez et al. 2018). We found that VIPP1 and VIPP2 proteins increased in GLA-treated cells, suggesting the activation of a chloroplast stress response (Fig. 1E). These results prompted us to investigate the effect of GLA on the photosynthetic light reactions. GLA decreased oxygen production and photosynthetic efficiency (Y(II)) at 50 μmol photons m^-2^ s^-1^ while having a mild effect on the *F*_v_/*F*_m_ ratio (Fig. 1F). In agreement with the blockage of the CBB cycle, GLA treatment led to an increase in NADPH fluorescence and oxygen consumption (Supplemental Fig. S3A and S3B).

We also analyzed the production of ROS following GLA treatment using the H_2_DCFDA probe as previously described (Perez-Perez et al. 2012a). Our results showed that GLA strongly increased the level of ROS within 30 min, remaining high for at least 8 h (Fig. 1G). To more specifically investigate the generation of H_2_O_2_ in GLA-treated cells, we used *Chlamydomonas* strains expressing the genetically encoded H_2_O_2_ sensor roGFP2-Tsa2ΔC_R_ in different cellular compartments (Niemeyer et al. 2021) (Supplemental Fig. S4). A rapid increase in H_2_O_2_ levels was detected in the cytosol and chloroplast compartments. However, chloroplast H_2_O_2_ levels decreased progressively over time, whereas cytosolic H_2_O_2_ remained elevated even after 8 h (Fig. 1H). Given that elevated ROS triggers autophagy in *Chlamydomonas* (Perez-Perez et al. 2012a, 2016; Mallén-Ponce and Pérez-Pérez 2023), we speculated that ROS generated by GLA treatment might contribute to the activation of this degradative process. To test this hypothesis, we analyzed ATG8 lipidation in *Chlamydomonas* cells treated with GLA and the ROS scavenger N-Acetyl-L-Cysteine (NAC). Our results revealed a decreased induction of autophagy by GLA in the presence of NAC (Fig. 1I), pointing to ROS generation as part of the cellular response that triggers autophagy in response to GLA. Taken together, our results strongly suggest that the inhibition of CO_2_ fixation by GLA leads to a pronounced chloroplast stress and enhances ROS production, which in turn triggers autophagy in *Chlamydomonas* cells.

### Deletion of phosphoribulokinase (PRK) triggers autophagy

To better characterize the connection between autophagy, carbon metabolism and ROS production, we investigated this catabolic process in the *Chlamydomonas prk* mutant, which lacks the PRK enzyme that catalyzes the conversion of ribulose-5-phosphate (Ru5P) to ribulose-1,5-bisphosphate (RuBP) (Fig. 1A). This mutant cannot grow in the absence of organic carbon and displays hypersensitivity to light (Boisset et al. 2023). Firstly, we performed a metabolomic analysis of WT and *prk* cells under medium light (50 μmol photons m^-2^ s^-1^) in the presence of acetate. Our results indicated that the *prk* mutant displayed a general metabolic imbalance since most of central carbon-derived metabolites were downregulated (Supplemental Fig. S5). Secondly, we investigated the regulation of autophagy in *prk* cells grown under different light and carbon regimes. For this, we monitored ATG8 and ATG8-PE levels in WT and *prk* cells grown mixotrophically in dark, low (20 μmol photons m^-2^ s^-1^) or medium (50 μmol photons m^-2^ s^-1^) light in exponential (0 h) and stationary (48 h) growth phase. Our data revealed that both ATG8 and ATG8-PE levels increased in the *prk* mutant in response to low or medium light but remained unaltered in dark conditions (Fig. 2A). An induction of autophagy under medium light was detected in *prk* cells at exponential growth phase, that further increased at stationary phase (Fig. 2A). To confirm that the activation of autophagy in the *prk* mutant was due to the absence of PRK, ATG8 lipidation was monitored in a complemented line. As expected, we found that the complemented strain 279-9 exhibited similar ATG8 levels to WT cells (Fig. 2B). Thirdly, we also studied the effect of carbon depletion on autophagy in the *prk* mutant. For this, WT and *prk* cells grown with acetate in low light were shifted to acetate-free medium. A gradual increase of ATG8 and ATG8-PE were detected in the *prk* mutant but not in WT cells (Fig. 2C). Finally, we tested the sensitivity of WT and *prk* strains to sublethal concentrations of the autophagic flux inhibitor ConcA. Interestingly, we found that the *prk* mutant was hypersensitive to autophagy inhibition under low or medium light but not in the dark (Fig. 2D), suggesting that autophagy is required for survival of *prk* cells under light stress.

**Figure 2.**
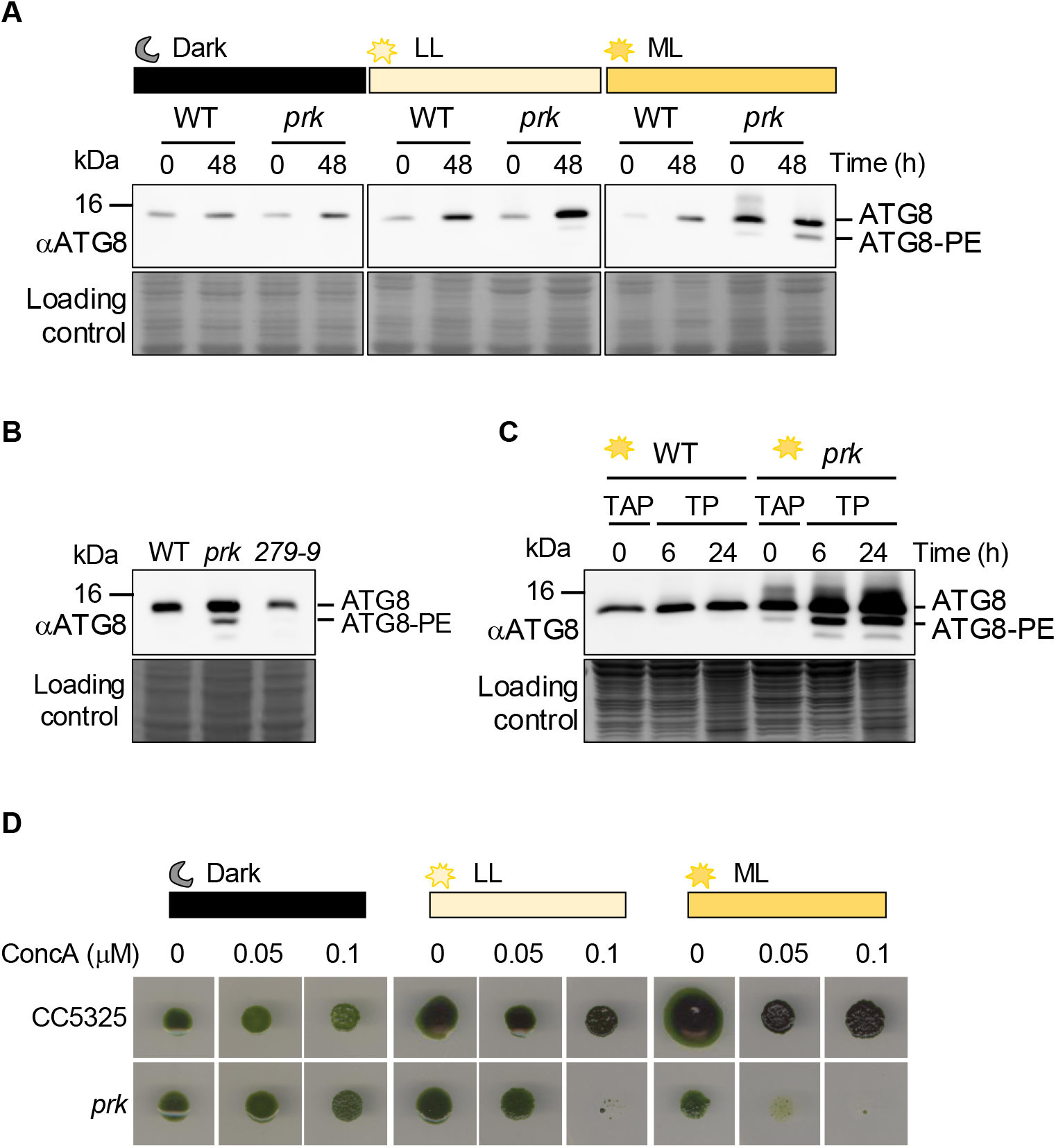
The *Chlamydomonas prk* mutant displays enhanced autophagy under standard growth conditions. **(A)** Immunoblot analysis of ATG8 protein levels in WT and *prk* mutant strains during the exponential (0 h) and stationary (48 h) growth phases in acetate-containing medium (TAP) under different light regimes: darkness, low light (LL; 20 μmol photons m^-2^ s^-1^), and medium light (ML; 50 μmol photons m^-2^ s^-1^). **(B)** Immunoblot analysis of ATG8 protein levels in WT, *prk* and the PRK-complemented line (279-9) grown in the presence of acetate in medium light. **(C)** Immunoblot analysis of ATG8 protein levels in WT and *prk* cells grown exponentially under mixotrophic conditions in TAP medium (0 h) and subsequently transferred to acetate-free medium (TP) for 6 or 24 h under medium light. **(D)** WT and *prk* cells grown exponentially in liquid TAP medium were subjected to 10-fold serial dilutions and spotted onto TAP agar plates containing 0, 0.05 or 0.1 μM concanamycin A (ConcA). Growth was documented after 7 days of incubation at 25°C in the dark or under continuous illumination (20 or 50 μmol photons m^-2^ s^-1^). Coomassie Brilliant Blue-stained gels were used as loading controls. Free ATG8 and lipidated ATG8 (ATG8-PE) are indicated on the right.

### ROS generation in the *prk* mutant under light and carbon stress

Given the tight connection between autophagy activation and oxidative stress in *Chlamydomonas* and plants (Perez-Perez et al. 2012b; Avin-Wittenberg 2019; Signorelli et al. 2019; Mallén-Ponce et al. 2023), we reasoned that the induction of this degradative process in the *prk* mutant might be linked to the production of ROS under light conditions. To explore this hypothesis, we determined the level of ROS in WT and *prk* cells grown in dark or different light intensities (20 or 50 μmol photons m^-2^ s^-1^), using the H_2_DCFDA probe as describe above. Our data revealed a high production of ROS in *prk* cells under low or medium light conditions (Fig. 3A). Furthermore, we analyzed the effect of the ROS scavenger NAC on autophagy activation in WT and *prk* cells grown in medium light. Our results indicated that the presence of NAC largely prevented the activation of autophagy in *prk* cells at stationary growth phase (Fig. 3B). Accordingly, a decreased level of ROS was detected in NAC-supplemented *prk* mutant cells (Fig. 3C).

**Figure 3.**
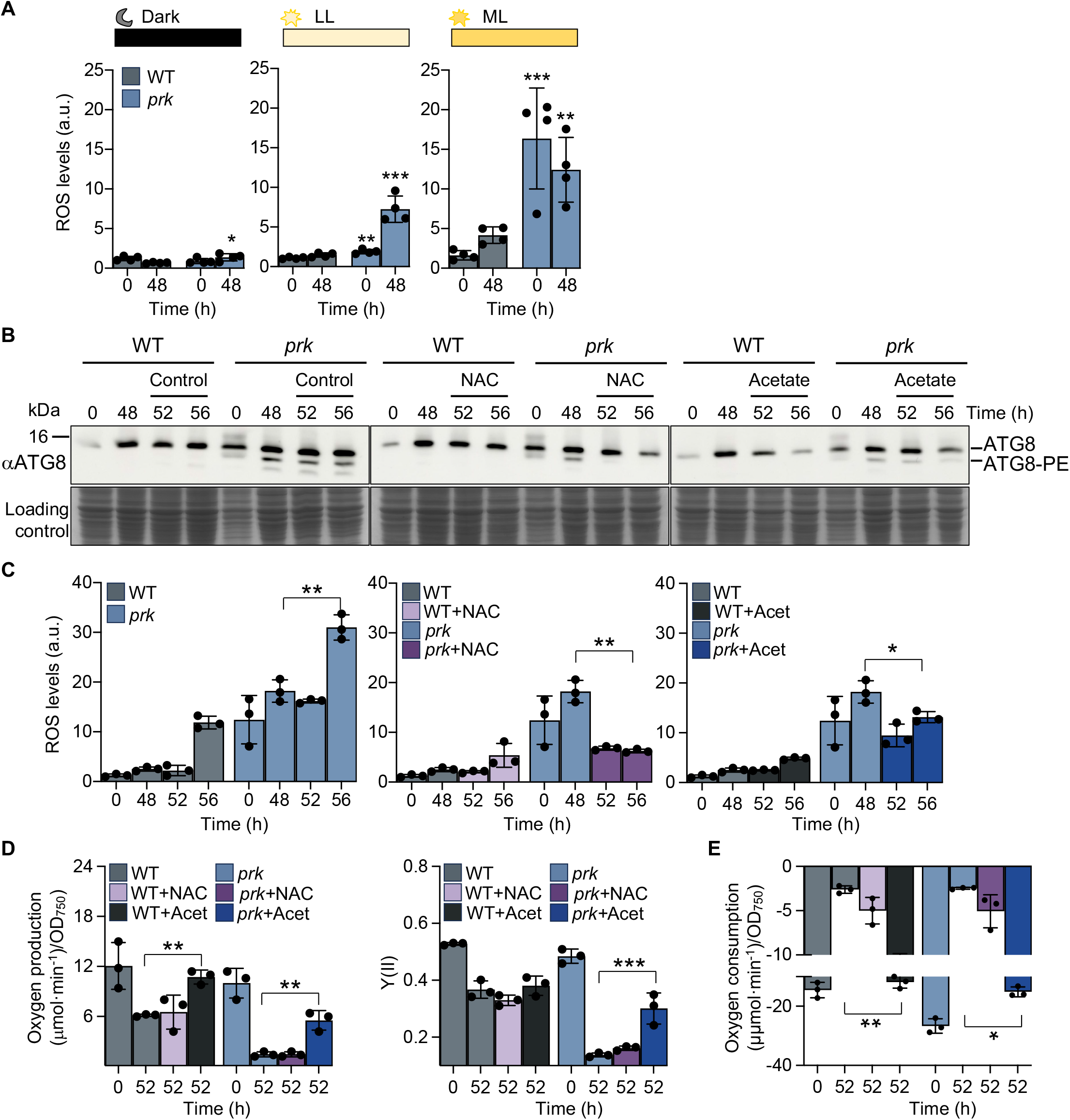
ROS associated to carbon deprivation activates autophagy in the *Chlamydomonas prk* mutant. **(A)** ROS levels in WT and *prk* cells during the exponential (0 h) and stationary (48 h) growth phases in acetate-containing medium under dark, low light (LL; 20 μmol photons m^-2^ s^-1^) or medium light (ML; 50 μmol photons m^-2^ s^-1^) conditions. **(B)** Immunoblot analysis of ATG8 protein levels in WT and *prk* strains. Cells were grown mixotrophically to the stationary phase (48 h) under medium light and subsequently treated with either 10 mM N-acetyl-L-cysteine (NAC, middel panel) or 15 mM acetate (right panel) for 4 h (52 h) or 8 h (56 h). Untreated cells were used as control (left panel). **(C)** ROS levels in WT and *prk* cells under the conditions described in *(B)*. **(D)** Oxygen evolution (left panel) and effective quantum yield of PSII (Y(II)) (right panel) in WT and *prk* cells grown under the conditions described in *(B)* in the absence or presence of NAC or acetate. **(E)** Oxygen consumption under the conditions described in *(B)*. Coomassie Brilliant Blue-stained gels were used as loading controls. Error bars represent SD. Values are expressed in arbitrary units (a.u.), with the untreated WT sample used as reference in (*A*, *C*, *D* and *E*). Asterisks indicate statistically significant differences determined by a two-tailed Student’s *t*-test: \*\*\**P*<0.001; \*\**P*<0.01; \**P*<0.05; and n.s. not significant ≥0.05. Data represent the mean of at least three independent biological experiments. Details of the statistical analyses are provided in Supplemental Table S3.

Since *prk* cells use acetate from the growth medium as carbon source, and *Chlamydomonas* quickly consumes acetate (Bogaert et al. 2019; Theis et al. 2019), we hypothesized that carbon limitation may participate in the activation of autophagy in this mutant. We thus determined the concentration of acetate after 24 and 48 h of growth in medium light. Acetate consumption was similar in both strains, being almost undetectable in the extracellular medium after 48 h (Supplemental Fig. S6A). We found that the addition of fresh acetate partially decreased the activation of autophagy by stationary growth in both WT and *prk* cells (Fig. 3B). Moreover, the production of ROS in the *prk* mutant partially decreased upon acetate supplementation (Fig. 3C), suggesting that the activation of autophagy in these cells might be linked to the generation of ROS due to a carbon starvation response.

Given that photosynthesis provides ATP and NADPH for carbon fixation, and the absence of PRK blocks the photosynthetic assimilation of CO_2_ (Boisset et al. 2023), we determined photosynthesis and respiration rates in WT and *prk* cells under exponential (0 h) and stationary (52 h) growth in medium light. Our results uncovered a decrease of oxygen production and photosynthetic yield Y(II) in WT cells at stationary growth phase that was more pronounced in the *prk* mutant (Fig. 3D). No significant effect was detected in *F*_v_/*F*_m_ ratio in both WT and *prk* cells (Supplemental Fig. S7). We also analyzed photosynthetic parameters in stationary WT and *prk* cells supplemented with NAC or acetate under the same light conditions. Whereas fresh acetate stimulated oxygen production and Y(II) in both WT and *prk* cells, NAC had no effect on these parameters (Fig. 3D). In close agreement with these results, we observed that oxygen consumption decreased similarly in both WT and *prk* strains at stationary growth phase while the addition of fresh acetate strongly stimulated respiration in these cells (Fig. 3E). Together, our results indicate that the inability of the *prk* mutant to assimilate CO_2_ resulted in decreased photosynthesis and the activation of ROS-linked autophagy in *Chlamydomonas*.

### Generation and characterization of a *Chlamydomonas* autophagy defective mutant by CRISPR-Cas9

To investigate the function of autophagy in the regulation of carbon metabolism in microalgae, we generated a *Chlamydomonas atg8* knockout mutant strain using CRISPR-Cas9 gene editing as previously described (Shin et al. 2016). To this aim, we introduced the sgRNA targeted to the first exon (E1) in the *ATG8* gene, the Cas9 protein and the pSL16 plasmid (containing the *APHVIII* paromomycin resistance gene) in WT cells by electroporation (Fig. 4A). The insertion of the *APHVIII* cassette in the *ATG8* gene was validated by PCR amplification of flanking sequences in paromomycin resistant clones. A DNA rearrangement in the E1 exon, but not in other regions of the *ATG8* gene, was detected in the *atg8* mutant (Supplemental Fig. S8A and S8B; Supplemental Table S2). Moreover, we performed qPCR and western blot analyses to show that the *atg8* mutant displayed negligible levels of *ATG8* mRNA and ATG8 protein, respectively (Fig. 4B and 4C; Supplemental Table S2). To further characterize the *atg8* mutant, we investigated autophagic flux by monitoring RPS6 and RPL37 protein abundance as previously described (Couso et al. 2018). For this, we analyzed RPS6 and RPL37 levels in WT and *atg8* cells, using as a negative control the cytosolic FKBP12 protein, whose abundance is independent of autophagy (Perez-Perez et al. 2010). We found that both ribosomal proteins accumulated to significantly higher levels in *atg8* cells compared to WT cells, indicating that autophagic flux was indeed impaired in this mutant (Fig. 4D). To rescue the *atg8* mutation, we generated an *atg8* complemented strain, named *atg8*::Strep-ATG8, expressing ATG8 tagged at the N terminus with the StrepII tag under the control of the *PSAD* promoter and terminator using the *Chlamydomonas* Modular Cloning (MoClo) Kit (Crozet et al. 2018; Niemeyer et al. 2021). Following the MoClo strategy, the StrepII-ATG8 part was assembled with a module containing the *APHVII* gene that confers resistance to hygromycin (Supplemental Fig. S8c). The resulting plasmid was used to transform *Chlamydomonas atg8* mutant cells and the expression of StrepII-tagged ATG8 was confirmed by western blot analysis with anti-CrATG8 antibodies (Supplemental Fig. S8D).

**Figure 4.**
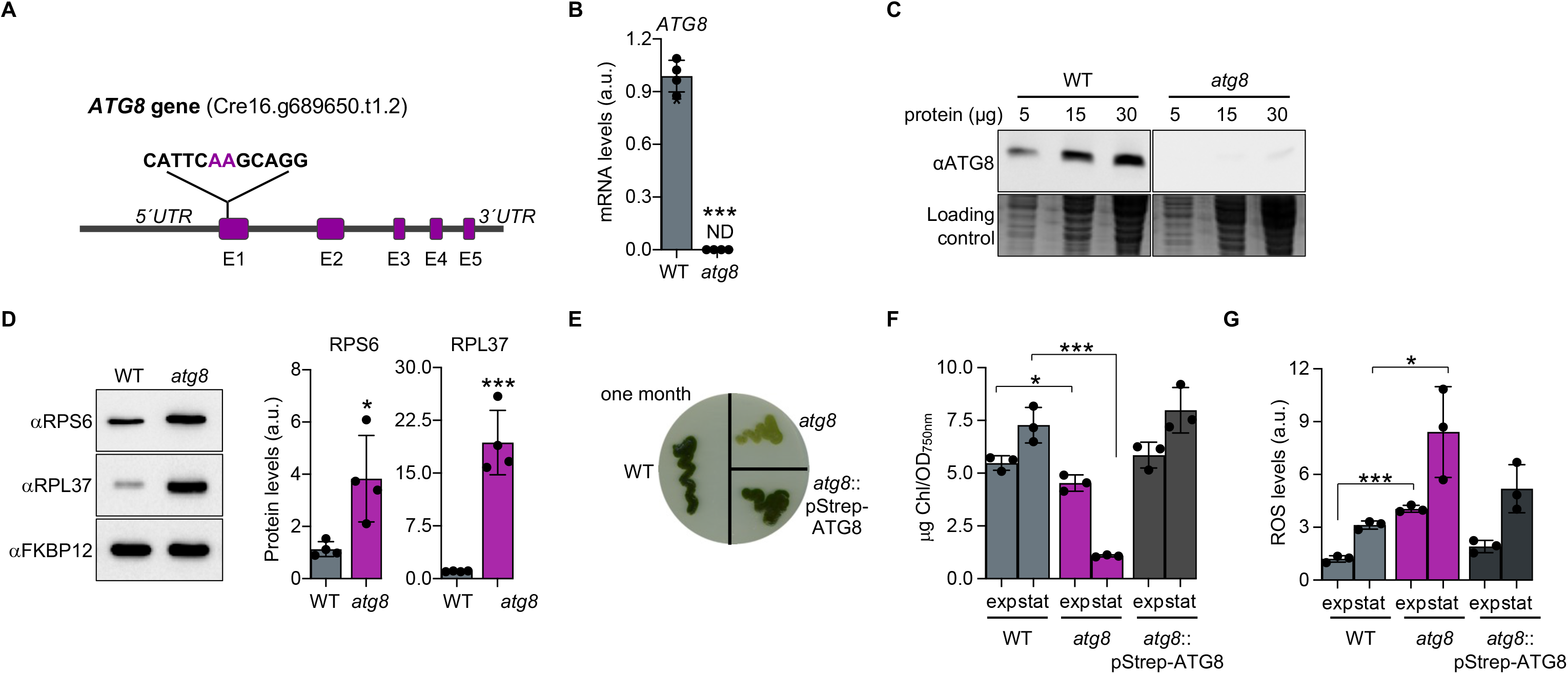
Generation and characterization of a *Chlamydomonas atg8* mutant lacking the core autophagy protein ATG8. **(A)** Schematic representation of the *ATG8* gene (Phytosome accession number: Cre16.g689650.t1.2) showing the five exons (E1-E5) and the 5’-UTR and 3’-UTR regions. The CRISPR/Cas 9 target sequence and the two nucleotides (AA; highlighted in magenta) recognized by the Cas9 nuclease are indicated. **(B)** qPCR analysis of *ATG8* mRNA abundance in WT and *atg8* mutant cells. Expression levels are shown in arbitrary units (a.u.), with the WT sample used as reference. Three biological replicates, each with three technical replicates, were analyzed per strain. ND, no signal detected. **(C)** Immunoblot analysis of increasing amounts of total protein extracts (5, 15 and 30 μg) from WT and *atg8* mutant cells. Endogenous ATG8 was detected using the anti-CrATG8 antibody. Coomassie Brilliant Blue-stained gels were used as loading control. **(D)** Autophagic flux assay in *atg8* mutant cells. Immunoblot analysis of proteins RPS6 and RPL37 ribosomal in exponentially growing WT and *atg8* mutant cells. 20 μg of total protein from each strain were probed with anti-CrRPS6 and anti-CrRPL37 antibodies, respectively. FKBP12 protein and Coomassie Brilliant Blue-stained gels were used as loading controls. Quantification of RPS6 (left panel) and RPL37 (right panel) protein levels is shown. **(E)** Growth phenotypes of WT, *atg8* and the complemented line *atg8*::Strep-ATG8 on solid medium under continuous illumination after 1 month of growth. **(F)** Chlorophyll contents of WT, *atg8* and *atg8*::Strep-ATG8 cells during the exponential (exp) and stationary (stat) growth phases in liquid TAP medium. **(G)** ROS levels in WT, *atg8* and *atg8*::Strep-ATG8 cells during the exponential (exp) and stationary (stat) growth phases as explained in *(F*). Error bars represent SD. Values are expressed in arbitrary units (a.u.), with WT cells used as the reference in (*D*) and WT in the exponential phase used as the reference in (*F*) and (*G*). Asterisks indicate statistically significant differences determined by a two-tailed Student’s *t*-test: \*\*\**P*<0.001; \*\**P*<0.01; \**P*<0.05; and n.s. not significant ≥0.05. Data represent the mean of at least three independent biological replicates. Details of the statistical analyses are provided in Supplemental Table S3.

Given that autophagy is activated in stationary growth in *Chlamydomonas* (Perez-Perez et al. 2010) or during senescence in plants (Yoshimoto et al. 2009), we analyzed the phenotype of the *atg8* mutant under prolonged growth. Consistent with previous observations in *Chlamydomonas* (Kajikawa et al. 2019), we observed that *atg8* mutant cells displayed a “yellowish” and senescent phenotype on solid medium at late stationary growth phase (Fig. 4E). Moreover, a significantly lower chlorophyll content was detected in the *atg8* mutant at stationary growth phase (Fig. 4F). To characterize further the phenotype of the *atg8* mutant, we determined the level of ROS, using the H_2_DCFDA probe, in exponential and stationary growth. This analysis revealed a higher ROS content in *atg8* mutant cells under both conditions (Fig. 4G).

### Autophagy is required for the recovery of *prk* mutant cells from carbon and light stress

To better explore the link between autophagy and carbon assimilation in *Chlamydomonas*, we analyzed the effect of sublethal concentrations of GLA (0.5 mM) in the *atg8* mutant. Strikingly, we found that the *atg8* mutant was hypersensitive to GLA under autotrophic conditions but not in the presence of acetate as an external carbon source (Fig. 5A). Consistent with this GLA-associated hypersensitivity, *atg8* cells exhibited a higher percentage of cell death and increased ROS production upon GLA treatment (Fig. 5B and 5C). Moreover, we carried out a comparative photosynthetic analysis of GLA-treated WT and *atg8* cells. Our results revealed that O_2_ evolution and Y(II) were further impaired in the *atg8* mutant in the presence of GLA (Fig. 5D). Likewise, the *F*_v_/*F*_m_ ratio and O_2_ consumption were significantly decreased in GLA-treated *atg8* cells, whereas no significant changes were detected in WT cells under the same condition (Fig. 5D).

**Figure 5.**
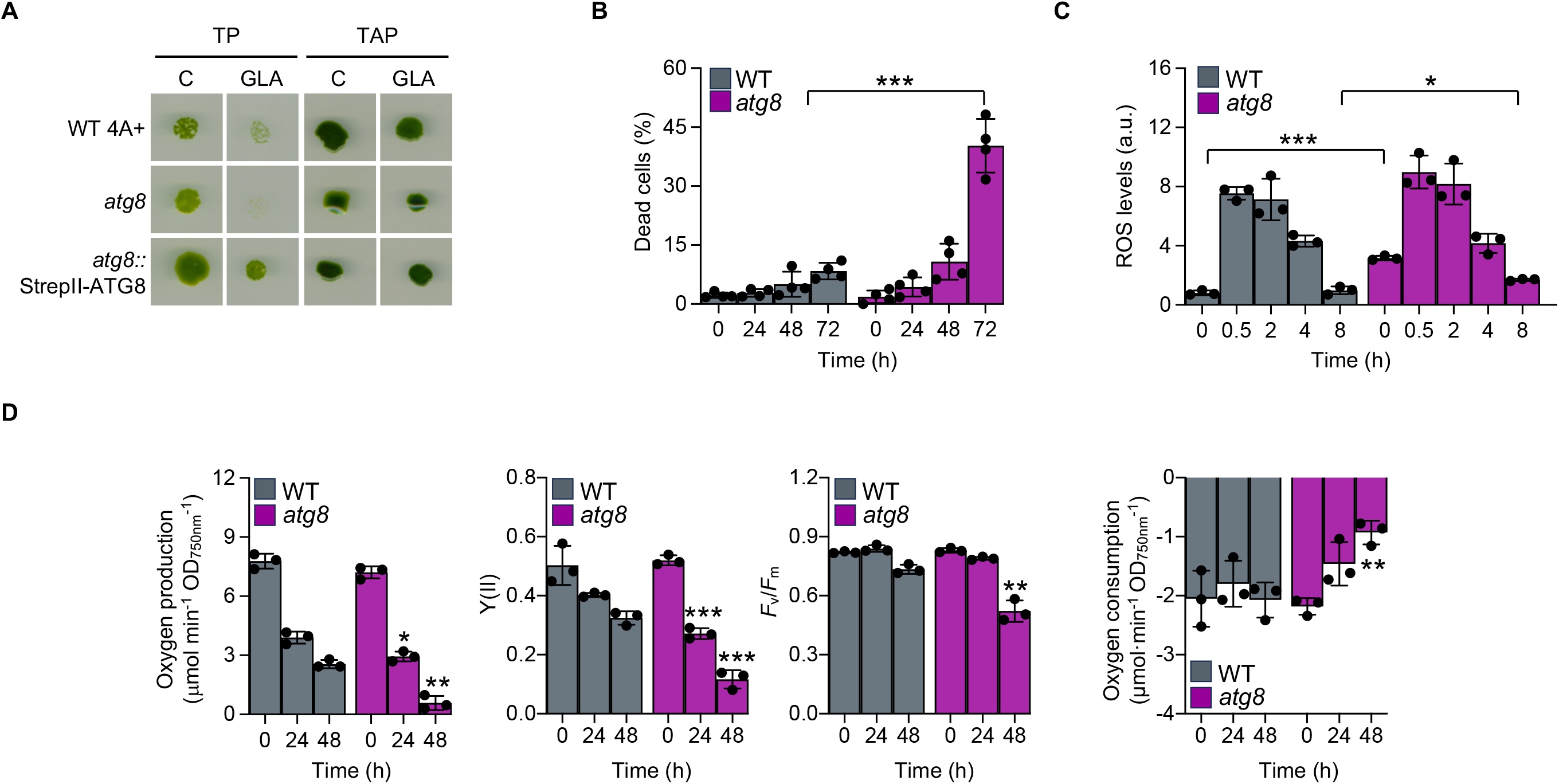
The *Chlamydomonas atg8* mutant exhibits hypersensitivity to carbon limitation caused by inhibition of CO_2_ fixation. **(A)** WT, *atg8* and *atg8*::Strep-ATG8 *Chlamydomonas* cells growing exponentially in acetate-free (TP) or acetate-containing (TAP) medium were subjected to 10-fold serial dilutions and spotted onto TP or TAP agar plates, respectively, supplemented with 0 (control, C) or 0.5 mM GLA. Plates were incubated at 25°C under continuous medium light conditions (50 μmol photons m^-2^ s^-1^) for 6-7 days. **(B)** Percentage of cell death in WT and *atg8* cells in the presence of GLA over time (0, 24, 48 and 72 h). **(C)** ROS levels in WT and *atg8* cells following GLA treatment for 0, 0.5, 2, 4 and 8 h. **(D)** Oxygen evolution, effective quantum yield of PSII (Y(II)), *F*_v_/*F*_m_ ratio and oxygen consumption in WT and *atg8* cells grown in the absence or presence of GLA. Error bars represent SD. Values are expressed in arbitrary units (a.u.), with the untreated WT sample used as reference in panels (*B*) and (*C*). Asterisks indicate statistically significant differences determined by a two-tailed Student’s *t*-test: \*\*\**P*<0.001; \*\**P*<0.01; \**P*<0.05; and n.s. not significant ≥0.05. Data represent the mean of at least three independent biological experiments. Details of the statistical analyses are provided in Supplemental Table S3.

Together, these findings strongly support a role of autophagy in carbon assimilation in *Chlamydomonas*. To test this hypothesis, we generated an *atg8 prk* double mutant by crossing the corresponding *atg8* and *prk* single-mutant strains (see Methods). The resulting progeny were screened by PCR, and the identity of the double mutant was subsequently confirmed by immunoblot analysis (Supplemental Fig. S9). We first characterized the response of *atg8 prk* and the corresponding single mutants to different light intensities. The WT, single-mutant and double-mutant strains exhibited similar growth under low light, although a growth defect was detected in the *prk* and the *atg8 prk* mutants under medium light (Fig. 6A). However, unlike WT and *atg8* strains, both the *prk* and *atg8 prk* mutants were unable to grow in high light (Fig. 6A). We therefore investigated the ability of the *prk* and *atg8 prk* strains to recover from high light stress. For this, cells growing under permissive low light conditions for 3 days, were then exposed to high light stress for 4 days, and finally transferred to medium light conditions for 1-day recovery period. Given the fast consumption of acetate from the medium (Supplemental Fig. S6A), *Chlamydomonas* cultures were supplemented with fresh acetate to prevent carbon limitation in the recovery assay. Our results show that the *prk* mutant resumed growth under medium light in acetate-supplemented cultures (Fig. 6B). However, in striking contrast, the *atg8 prk* double mutant was unable to restore growth following high light exposure (Fig. 6B), indicating that autophagy is required for the recovery of *prk* cells from light stress.

**Figure 6.**
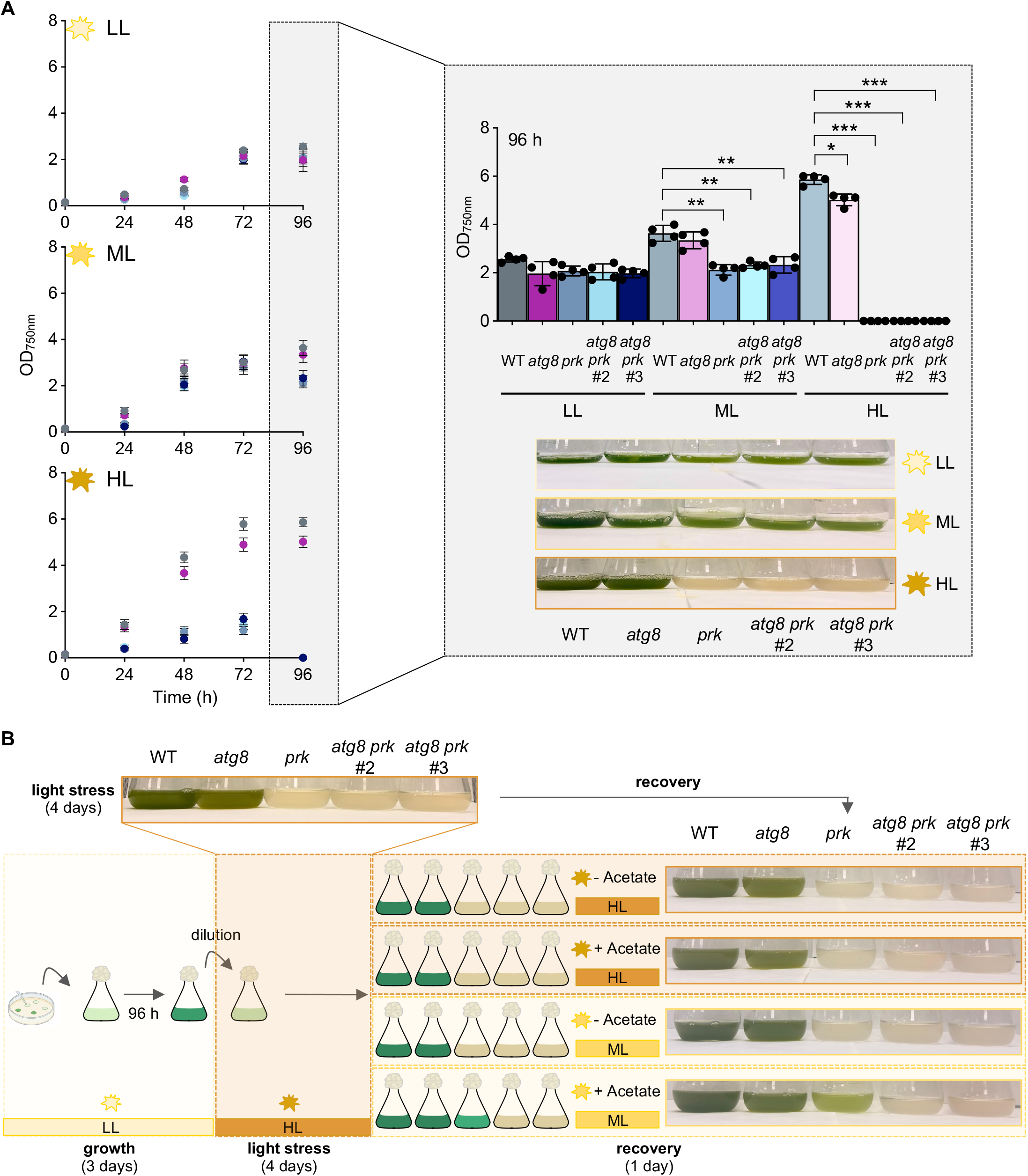
The double *Chlamydomonas atg8 prk* mutant is impaired during the recovery after high light stress. **(A)** Growth curves of WT, *atg8*, *prk*, *atg8 prk #2* and *atg8 prk #3* cells grown in acetate-containing (TAP) medium under low light (LL; 20 μmol photons m^-2^ s^-1^, upper panel), medium light (ML; 50 μmol photons m^-2^ s^-1^, upper panel) and high light (HL; 350 μmol photons m^-2^ s^-1^, lower panel). The right panel shows a graphical representation of growth (upper panel) and representative images (lower panel) of the different strains after 96 h of growth for 96 h under LL, ML and HL. **(B)** Schematic representation of the recovery experiment. Briefly, *Chlamydomonas* cells grown in TAP medium grown in LL conditions for 3 days were diluted into fresh TAP medium to an initial OD_750nm_ of 0.15 and exposed to HL for 4 days. Following HL stress, cells were transferred to ML conditions for 1 day either in the absence or presence of fresh 15 mM acetate. In parallel, cells were maintained under high light conditions for an additional day in either the absence or presence of fresh acetate. Representative images of the five strains after high light stress and during the recovery phase are shown. Asterisks indicate statistically significant differences determined by a two-tailed Student’s *t*-test: \*\*\**P*<0.001; \*\**P*<0.01; \**P*<0.05; and n.s. not significant ≥0.05. Data represent the mean of at least three independent biological experiments. Details of the statistical analyses are provided in Supplemental Table S3.

## Discussion

Autophagy is increasingly recognized as an important component of the metabolic and stress-response networks that coordinate nutrient utilization, cellular homeostasis and environmental acclimation in photosynthetic organisms. Although autophagy induction has been extensively documented in plants and microalgae, the relationship between photosynthetic carbon assimilation and autophagy regulation remains less well understood. Our results demonstrate that impairment of CBB cycle activity promotes autophagy activation in *Chlamydomonas* and suggest that this response is closely associated with ROS accumulation arising from an imbalance between photosynthetic electron transport and carbon fixation.

The results obtained using both GLA and the *prk* mutant indicate that disruption of CBB cycle activity is associated with autophagy activation in *Chlamydomonas*. Despite differences in experimental design and timescale, both approaches converge on a common metabolic consequence: the inability to efficiently regenerate RuBP and sustain photosynthetic CO₂ fixation. Autophagy increased further when carbon availability became more restrictive, following either from acetate depletion during stationary growth or from transfer of *prk* cells to acetate-free medium. Thus, autophagy appears to respond not only to carbon limitation itself but also to restrictions in photosynthetic carbon assimilation. This interpretation is consistent with studies in plants showing that autophagy is activated under conditions that limit carbon or energy availability, including extended darkness, reduced carbohydrate availability and impaired energy homeostasis (Thompson et al. 2005; Izumi et al. 2013; Avin-Wittenberg et al. 2015; Soto-Burgos and Bassham 2017). Accordingly, autophagy-deficient mutants exhibit alterations in carbon metabolism and reduced fitness under carbon-limiting conditions (Guiboileau et al. 2013; Avin-Wittenberg et al. 2015; Barros et al. 2017; McLoughlin et al. 2020). In *Chlamydomonas*, genetic disruption of ATG8 similarly affects carbon storage and remobilization under nutrient-limiting conditions (Kajikawa et al. 2019). The light dependence of autophagy activation by both GLA and loss of PRK further suggests that the relevant signal arises from the interaction between photosynthetic activity and defective carbon fixation rather than from carbon limitation alone.

Restriction of CBB cycle activity while photosynthetic electron transport remains active can lead to excess reducing power and ROS production, as described in higher plants when carbon assimilation becomes limiting (Foyer and Noctor 2005; Mittler 2017). The photosynthetic and chloroplast stress responses observed following PRK inhibition in *Chlamydomonas* are consistent with such a redox imbalance. Accordingly, ROS accumulated in both GLA-treated and *prk* cells under illumination, whereas NAC attenuated autophagy activation. These findings are consistent with the established role of ROS in autophagy regulation in plants and microalgae (Xiong et al. 2007; Perez-Perez et al. 2012a, 2016; Signorelli et al. 2019; Mallén-Ponce and Pérez-Pérez 2023) and with chloroplast ROS acting as retrograde signals during photosynthetic stress in plants (Exposito-Rodriguez et al. 2017; Foyer and Hanke 2022).

A connection between chloroplast dysfunction, ROS and autophagy is also emerging from studies in *Chlamydomonas*. Perturbation of chloroplast lipid metabolism, protein import or proteostasis induces autophagy and cellular responses associated with chloroplast quality control (Ramundo et al. 2014, 2020; Heredia-Martinez et al. 2018; Xing et al. 2022). More recently, Yang et al. (Yang et al. 2026) showed that disruption of chloroplast protein import results in chloroplast dysfunction associated with ROS accumulation, autophagy and extensive metabolic and organellar remodeling. These studies and the rapid chloroplast oxidation and sustained cytosolic oxidative response observed following GLA treatment support a model in which disruption of carbon fixation alters chloroplast redox homeostasis, generating signals that contribute to autophagy activation.

The characterization of the CRISPR-Cas9-generated *atg8* mutant provides insight into the physiological function of this response. Kajikawa et al. (Kajikawa et al. 2019) previously described an insertional *atg8* mutant exhibiting reduced survival and accelerated chlorophyll loss under nitrogen, sulfur and phosphate deficiency and showed that ATG8 contributes to starch accumulation during phosphate limitation and lipid remobilization following nitrogen resupply. The lower chlorophyll content and senescence-like phenotype of our CRISPR-derived *atg8* mutant are consistent with these findings and support a role for autophagy in maintaining cellular fitness during prolonged metabolic stress. Our results extend this function to redox homeostasis, as *atg8* cells accumulated higher ROS levels during both exponential and stationary growth. This phenotype is consistent with studies in plants showing that autophagy-deficient mutants accumulate oxidative damage and undergo premature senescence (Xiong et al. 2007; Yoshimoto et al. 2009; Guiboileau et al. 2013; Masclaux-Daubresse et al. 2014). Selective autophagy also contributes to chloroplast quality control and removal of damaged chloroplast material under stress (Izumi et al. 2017; Woodson 2019; Otegui et al. 2024). Thus, nutrient and carbon recycling may operate together with cellular quality control to explain the contribution of autophagy to metabolic and redox homeostasis in photosynthetic cells.

The enhanced sensitivity of *atg8* cells to GLA under autotrophic conditions further supports a protective function for autophagy when photosynthetic carbon assimilation is compromised (Fig. 5). Loss of ATG8 was associated with enhanced ROS accumulation, cell death and impairment of photosynthetic performance following PRK inhibition, suggesting that autophagy limits the downstream consequences of the resulting metabolic and redox imbalance, by faciliting the removal and recycling of damaged cellular components. Analysis of the *atg8 prk* double mutant extends this protective role to recovery from light-induced metabolic stress. Although both *prk* and *atg8 prk* mutants were unable to grow under high light, only *prk* cells resumed growth after transfer to permissive conditions in the presence of fresh acetate (Fig. 6). Thus, autophagy does not appear to only compensate directly for defective carbon fixation during sustained high light but contributes to restoring cellular homeostasis once the stress is relieved. Recovery is likely to require metabolic reorganization together with the removal or recycling of components damaged during stress. In plants, autophagy contributes to chloroplast quality control and turnover under photooxidative conditions (Izumi et al. 2017; Woodson 2019; Otegui et al. 2024). Although the autophagic substrates involved in *prk* recovery remain unknown, the inability of *atg8 prk* cells to resume growth is consistent with a requirement for autophagic recycling during recovery from prolonged impairment of photosynthetic carbon assimilation.

Overall, our findings support a model in which carbon assimilation, redox regulation and autophagy are interconnected in *Chlamydomonas*. Impairment of CO₂ fixation generates an imbalance between light-driven electron transport and carbon metabolism, promoting chloroplast stress and ROS production. These signals contribute to autophagy activation, which in turn limits stress-associated damage and facilitates recovery when favorable conditions are restored. Together with previous studies in microalgae and plants, our results support autophagy as an integral component of the mechanisms coordinating carbon metabolism, photosynthetic performance and redox homeostasis in photosynthetic eukaryotes.

## Author contributions

MEPP designed and supervised the study, with input from all co-authors. SGA, MJMP, IMP, LHM, AMQM and MEPP performed the majority of the experiments. SGA, MJMP, LHM, JLC and MEPP designed the experimental set up and analyzed the results. PC and SL contributed to the characterization of the *prk* mutant; MS assisted with the ROS compartmentalization experiments and the analysis of the chloroplast stress response; and CR designed the *atg8 prk* double mutant and contributed to its characterization. MEPP wrote the manuscript, with contributions and feedback from all co-authors.

## Acknowledgments

We thank C. Parejo for technical assistance with metabolite analyses at the IBVF Chromatography Facility and J.E. Frías for technical support at the IBVF Biological Culture Facility.

## Data availability

The data underlying this article are available in the article and in its online supplementary material.

