## Supplemental material for "Inhibition of carbon fixation activates ROS-linked autophagy in the single-celled green alga *Chlamydomonas reinhardtii*"

**Supplemental Information**

**Supplemental Figure S1. Effects of Calvin-Benson-Bassham (CBB) cycle inhibition in *Chlamydomonas*. (A)** WT *Chlamydomonas* cells growing exponentially in acetate-free (TP) or acetate-containing (TAP) medium were subjected to 10-fold serial dilutions and spotted onto TP or TAP agar plates supplemented with 0 or 0.5 mM glycolaldehyde (GLA). Plates were incubated at 25ºC under continuous medium-light conditions (50 µmol photons $m^{-2}$ $s^{-1}$) for 6-7 days. **(B)** Relative abundance of CBB cycle intermediates in WT *Chlamydomonas* cells treated with 1 mM GLA. Ru5P, ribulose-5-phosphate; Xu5P, xylulose-5-phosphate. Error bars represent SD from three independent biological replicates. Asterisks indicate statistically significant differences as determined by one-way ANOVA followed by Bonferroni's test: ****P*<0.001; ***P*<0.01; **P*<0.05; ns, not significant, ≥0.05. Details of the statistical analyses are provided in Supplemental Table S4.

**Supplemental Figure S2. Effects of glycolaldehyde (GLA) on autophagy activation.** Immunoblot analysis of ATG8 protein levels in WT *Chlamydomonas* cells grown under autotrophic conditions and treated with increasing concentrations of GLA (0, 0.5, 1 and 2 mM) for 24 h. Coomassie Brilliant Blue-stained gel was used as loading control.

**Supplemental Figure S3. Photosynthetic parameters in cells subjected to Calvin-Benson-Bassham (CBB) inhibition. (A)** NADPH fluorescence in WT cells before treatment (0 h) and after treatment with 1 mM glycolaldehyde (GLA) for 0.5, 1, 2, 4 and 8 h under medium light conditions (50 µmol photons $m^{-2}$ $s^{-1}$). **(B)** Oxygen consumption in WT cells under the conditions described in *(A)*. Error bars represent SD. Values are expressed in arbitrary units (a.u.), with the untreated WT sample used as the reference. Asterisks indicate statistically significant differences as determined by one-way ANOVA followed by Bonferroni´s test: ****P*<0.001; ***P*<0.01; **P*<0.05; and not significant ≥0.05. Data represent the mean of four independent biological experiments. Details of the statistical analyses are provided in Supplemental Table S4.

**Supplemental Figure S4. Schematic representation of roGFP2-Tsa2ΔCR $H_2O_2$ sensors targeted to different subcellular compartments in *Chlamydomonas*. (A)** Schematic representation of the genetic constructs used to target the roGFP2-Tsa2ΔCR $H_2O_2$ sensor to the chloroplast stroma or cytosol. The constructs contain the spectinomycin resistance cassette (*Sp*$^R$; *aadA* resistance marker); the *HSP70A-RBCS2* promoter (P*AR*); the *CDJ1* (light green), or *PAR* (gray) targeting sequences for localization of the sensor to the stroma or cytosol, respectively; *roGFP-Tsa2ΔCR* coding sequence; a *STOP* codon; and the *RPL23* terminator (t*RPL23*). Adapted from Niemeyer et al. (2021). **(B)** Schematic representation of a *Chlamydomonas* cell illustrating the putative accumulation of reactive species oxygen (ROS) in different subcellular compartments (chloroplast stroma and cytosol) under stress conditions.

**Supplemental Figure S5. Metabolomic analysis of WT and *prk* mutant *Chlamydomonas* cells.** Relative abundance of Calvin-Benson-Bassham (CBB) cycle intermediates **(A)**; NADPH **(B)**; central carbon metabolites **(C)**; and tricarboxylic acid (TCA) cycle intermediates **(D)**. WT and *prk* cells were grown mixotrophically and harvested during the exponential (0 h) and stationary (48 h) growth phase. Ru5P, ribulose-5-phosphate; Xu5P, xylulose-5-phosphate; RuBP, ribulose-1,5-bisphosphate; 3PGA, 3-phosphoglycerate; G3P, glyceraldehyde-3-phosphate; E4P, Erythrose-4-phosphate; R5P: Ribose-5-phosphate; DHAP: Dihydroxyacetone phosphate; PEP, phosphoenolpyruvate; Pyr, pyruvate; Acetyl-CoA, acetyl coenzyme A. Error bars represent SD from four independent biological replicates. Asterisks indicate statistically significant differences determined by a two-tailed Student's *t*-test: ***$P$<0.001; **$P$<0.01; *$P$<0.05; and n.s. not significant ≥0.05. Details of the statistical analyses are provided in Supplemental Table S4.

**Supplemental Figure S6. Acetate content in WT and *prk* mutant *Chlamydomonas* cells.** Acetate amount in the medium from WT and *prk* strains grown in TAP medium for 0, 24 and 48 h under medium light conditions (ML, 50 μmol photons $m^{-2}$ $s^{-1}$). Error bars represent SD. Asterisks indicate statistically significant differences as determined by one-way ANOVA followed by Bonferroni´s test: ***$P$<0.001; **$P$<0.01; *$P$<0.05; ns, not significant ≥0.05. Data represent the mean of seven independent biological experiments. Details of the statistical analyses are provided in Supplemental Table S4.

**Supplemental Figure S7. Maximum quantum efficiency of PSII ($F_v/F_m$) in WT and *prk* mutant *Chlamydomonas* cells.** WT and *prk* cells were grown under medium light conditions (ML, 50 μmol photons $m^{-2}$ $s^{-1}$) and analyzed during the exponential (0 h) and stationary (48 h) growth phases. The maximum quantum efficiency of PSII ($F_v/F_m$) was determined for each condition. Error bars represent SD. Asterisks indicate statistically significant differences as determined by one-way ANOVA followed by Bonferroni´s test: ***$P<0.001$; **$P<0.01$; *$P<0.05$; ns, not significant ≥0.05. Data represent the mean of three independent biological experiments. Details of the statistical analyses are provided in Supplemental Table S4.

**Supplemental Figure S8. Schematic representation of the *atg8* and the complemented *atg8*::Strep-tagged ATG8 lines. (A)** *ATG8* gene (Cre16.g689650.t1.2) showing its five exons (E1-5) and the 5´UTR and 3´UTR regions. The target sequence and the two nucleotides (AA, highlighted in magenta) recognized by the Cas9 protease where paromomycin cassette is inserted are shown. The primer pairs (a/a´, b/b´, c/c´ and d/d´; listed in Supplemental Table S2) used to check the paromomycin insertion by PCR tests are indicated. The arrow indicates the primer sense. **(B)** Insertional mutagenesis PCR tests using *ATG8* primers (a/a´, c/c´ and d/d´) and *APHVIII* primers (b/b´) in WT and *atg8* cells. The DNA fragment sizes are indicated on the left. **(C)** Schematic representation of the different parts of the StrepII-tagged ATG8 construct: hygromycin ($Hyg^R$) resistance cassette (*APHIV* resistance marker); *PSAD* promotor (P*PSAD*); *StrepII* sequence, *ATG8* codon sequence; a STOP codon; and the *PSAD* terminator (t*PSAD*). **(D)** Immunoblot analysis of ATG8 protein from WT, *atg8* and *atg8*::Strep-ATG8 cells growing mixotrophically at the exponential phase. Immunoblot analysis of FKBP12 was used as loading control. ATG8 and Strep-tagged ATG8 are indicated on the right.

**Supplemental Figure S9. Generation of *atg8 prk* double mutants.** Immunoblot analysis using anti-CrATG8 (upper panel) and anti-CrPRK (lower panel) antibodies to confirm the double mutation in *atg8 prk #2* and *atg8 prk #3* cells. Coomassie Brilliant Blue-stained gels were used as loading control.

**Figure Suppl. S1**

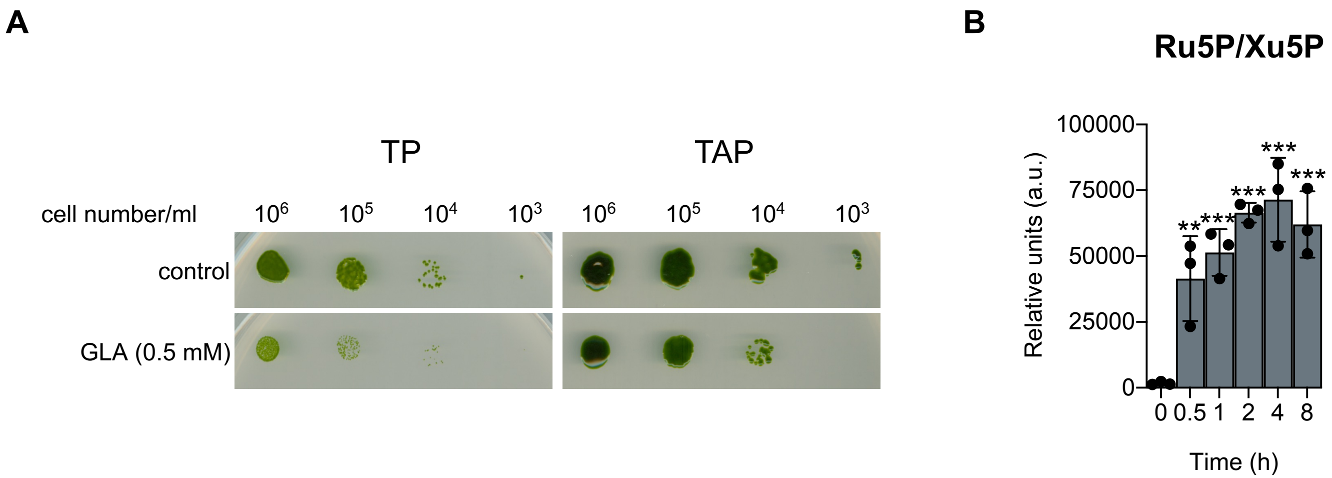

**Figure Suppl. S2**

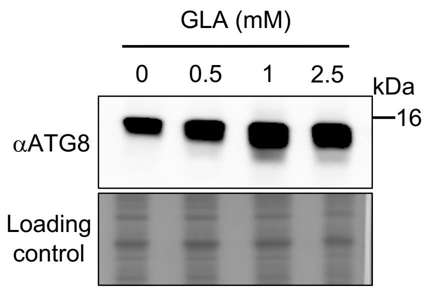

**Figure Suppl. S3**

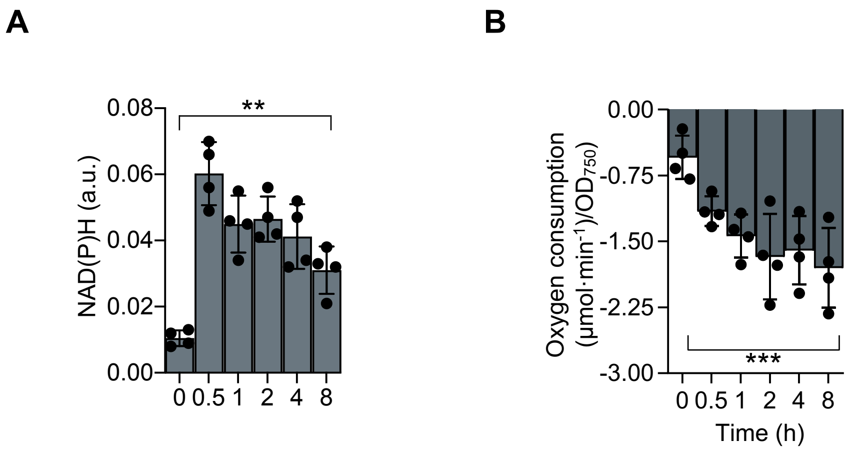

**Figure Suppl. S4**

A

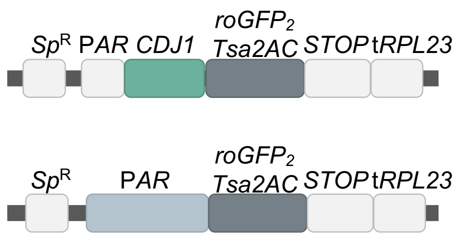


B

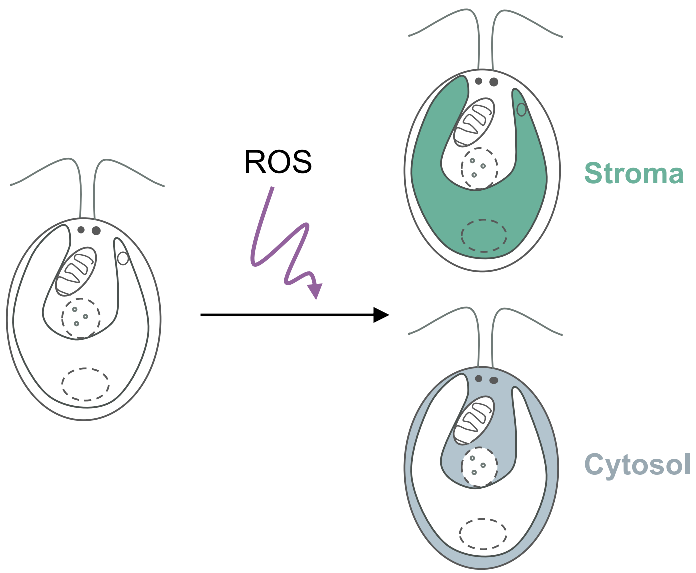

**Figure Suppl. S5**

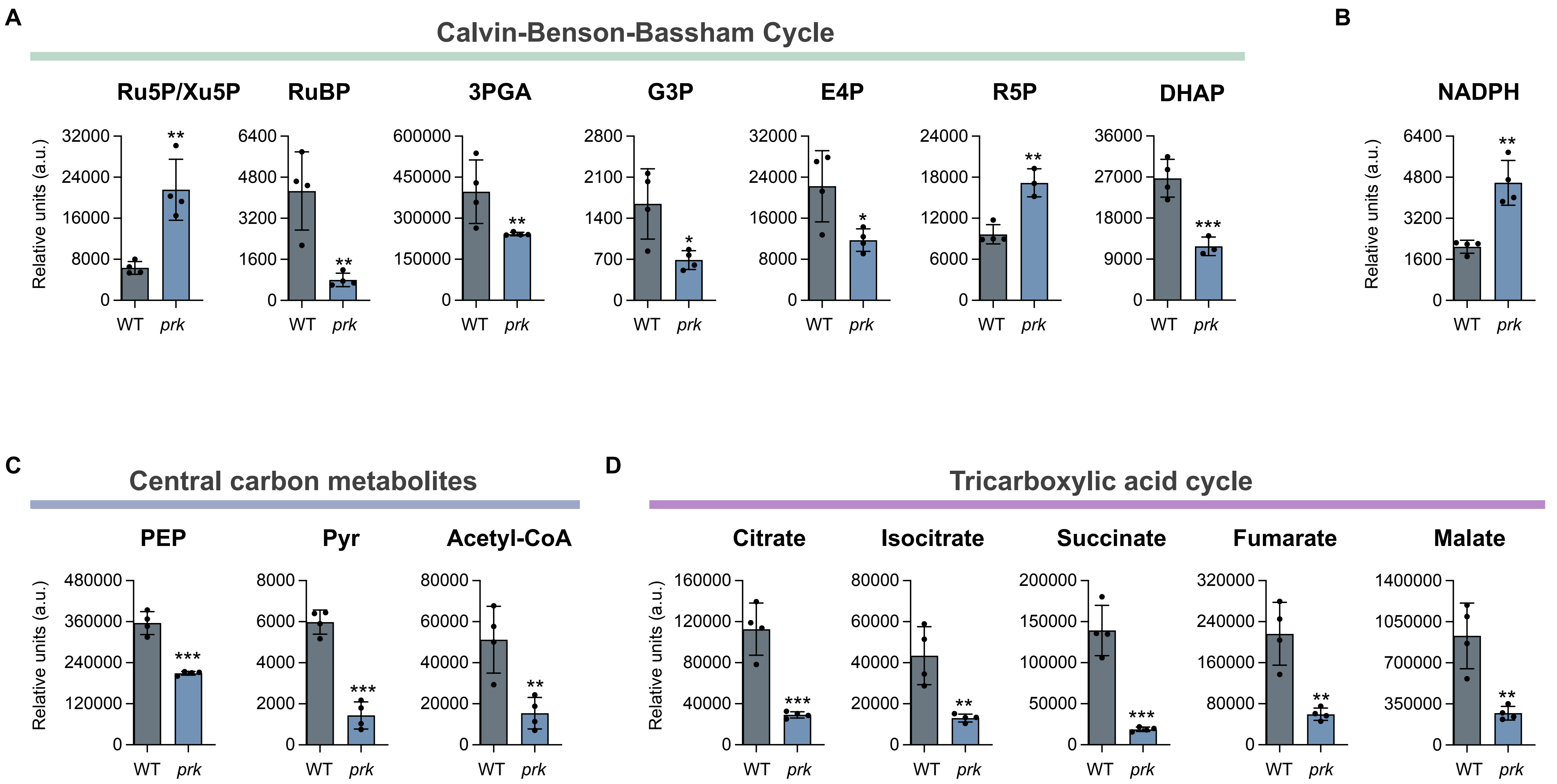

**Figure Suppl. S6**

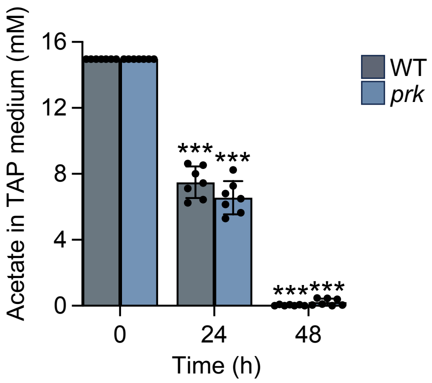

**Figure Suppl. 7**

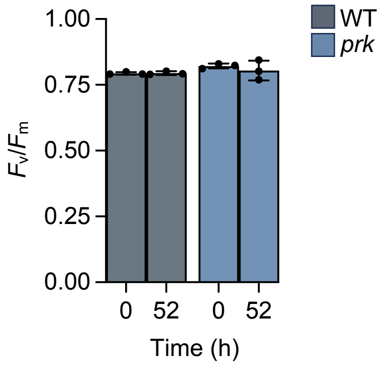

**Figure Suppl. S8**

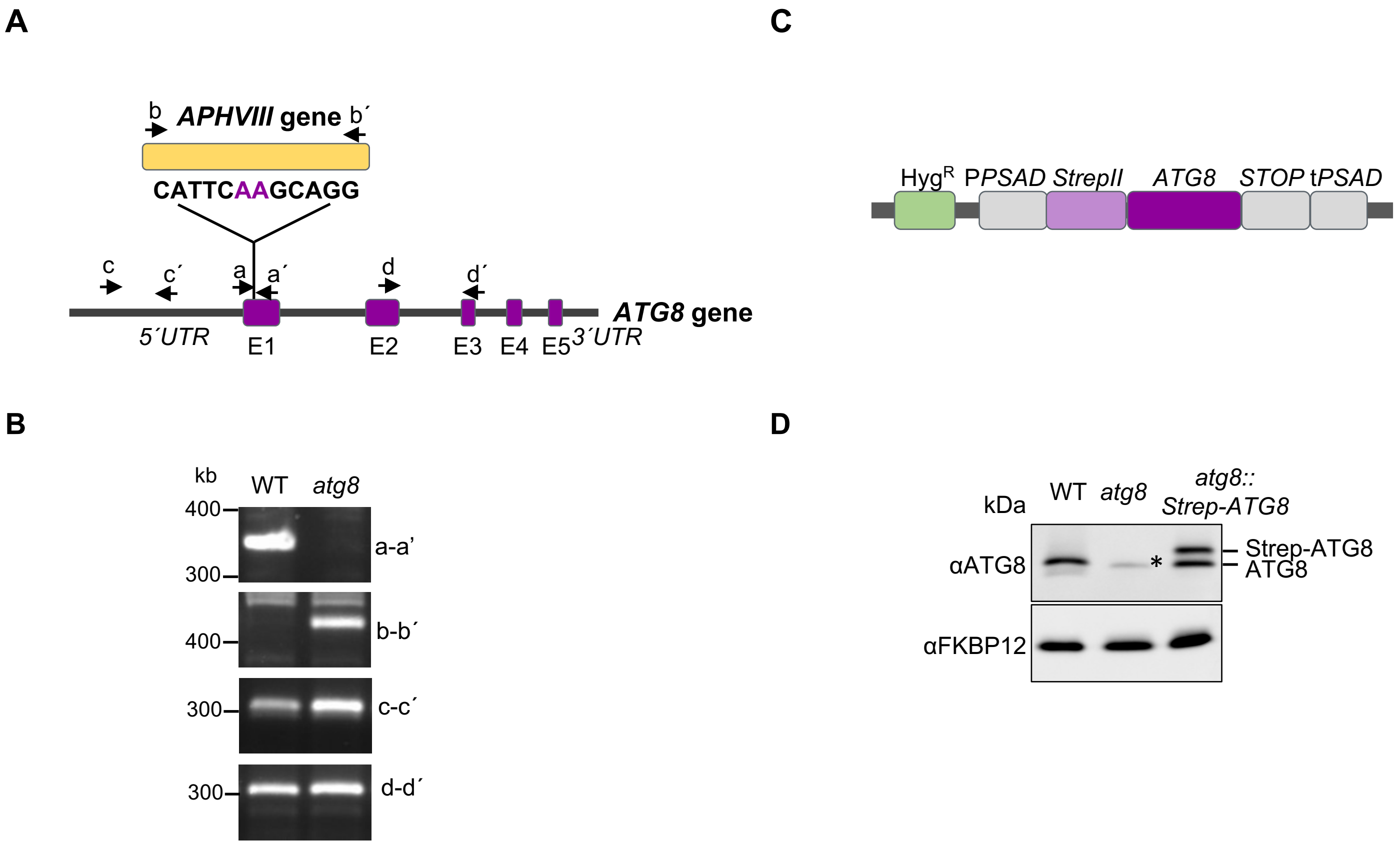

**Figure Suppl. S9**

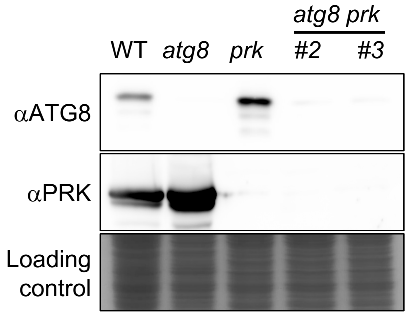

**Supplemental Table S1.** Chlamydomonas strains used in this work.

| Strains | Main characteristics | Reference |
|---|---|---|
| **WT4A+ (CC-4051)** | Wild-type *Chlamydomonas* strain. | Chlamydomonas Resource Center (http://www.chlamycollection.org). |
| ***atg8*** | *Chlamydomonas atg8* mutant strain generated by CRISPR-Cas9. The *ATG8* gene is disrupted by the insertion of the *APHVIII* cassette in the first exon. This strain is paromomycin-resistant. | This study. |
| ***atg8*::StrepII-ATG8** | *Chlamydomonas* ATG8-complemented line. The StrepII-tagged *ATG8* gene is expressed in the *Chlamydomonas atg8* mutant under the control of the *PSAD* promoter and terminator. This strain is hygromycin-resistant. | This study. |
| **WT (CC-4533)** | Wild-type *Chlamydomonas* strain. | Clip Library. Chlamydomonas Resource Center (http://www.chlamycollection.org). Li et al. (2019). |
| ***prk*** | *Chlamydomonas prk* mutant strain. The PRK gene is disrupted by the insertion of *APHVIII* gene in the exon 7. This strain is paromomycin-resistant. | Clip Library. Chlamydomonas Resource Center (http://www.chlamycollection.org). Li et al. (2019); Boisset et al. (2023). |
| ***atg8 prk*** | *Chlamydomonas atg8 prk* double mutant strain generated by genetic crosses between the single *atg8* and *prk* mutants. This strain is paromomycin-resistant. | This study. |
| **UVM4+** | *Chlamydomonas* strain derived from the cell wall-deficient cw15-302 background. The strain is optimized for high-level nuclear transgene expression and displays reduced gene silencing. | Neupert et al. (2009). |
| **ARpro** | A UVM4+-derived strain expressing the genetically encoded $H_2O_2$ sensor roGFP2-Tsa2ΔCR in the cytosol. | Niemeyer et al. (2021). |
| **CDJ1** | A UVM4+-derived strain expressing the genetically encoded $H_2O_2$ sensor roGFP2-Tsa2ΔCR in the chloroplast stroma. | Niemeyer et al. (2021). |


Li X, Patena W, Fauser F, Jinkerson RE, Saroussi S, Meyer MT, Ivanova N, Robertson JM, Yue R, Zhang R, Vilarrasa-Blasi J, Wittkopp TM, Ramundo S, Blum SR, Goh A, Laudon M, Srikumar T, Lefebvre PA, Grossman AR, Jonikas MC (2019). ***Nature Genetics*** 51(4):627-635.

Boisset ND, Favoino G, Meloni M, Jomat L, Cassier-Chauvat C, Zaffagnini M, Lemaire SD, Crozet P (2023). ***Front Plant Sci.*** 14:1230723.

Neupert, J, Karcher D, Bock R (2009). ***The Plant Journal*** 57(6):1140-1150.

Niemeyer J, Scheuring D, Oestreicher J, Morgan B, Schroda M (2021). ***The Plant Cell*** 33(9):2935-2949.

**Supplemental Table S2.** Primer sequences used for the characterization of *Chlamydomonas atg8* mutant.

| Primer name | Sequence (5'-3') | Characteristics | Reference |
|---|---|---|---|
| ***a***<br>***a´*** | 5´-CGTTGCCCATAGAACTGCC-3´<br>5´-CCGGCACTCACTGGAATTC-3´ | Forward primer located 150 bp upstream of the Cas9 cleavage site.<br>Reverse primer located 150 bp downstream of the Cas9 cleavage site. | This study. |
| ***b***<br>***b´*** | 5´-CAAACCGCCAGATCCTCGTC-3´<br>5´-GTTTGTCAAGGTGGCAGCTC-3´ | Forward primer located within the *APHVIII* gene.<br>Reverse primer located within the *APHVIII* gene. | This study. |
| ***c***<br>***c´*** | 5´-GTGGTCAGCGGGTATTTGG-3´<br>5´-GGCAGTTCTATGGGCAACG-3´ | Forward primer located upstream of the Cas9 cleavage site.<br>Reverse primer located downstream of the Cas9 cleavage site. | This study. |
| ***d***<br>***d´*** | 5´-GAATTCCAGTGAGTGCCGG-3´<br>5´-GAGGCACGGCTAGCACTAG-3´ | Forward primer located upstream of the Cas9 cleavage site.<br>Reverse primer located downstream of the Cas9 cleavage site. | This study. |
| ***5´ATG8*** | 5´-CTCGGAATACGCCAGCTTTT-3´ | Forward primer for *ATG8* gene expression analysis by RT-qPCR. | Pérez-Martín et al. (2014) |
| ***3´ATG8*** | 5´-TTCGGTCTGGGTACTTCTCC-3´ | Reverse primer for *ATG8* gene expression analysis by RT-qPCR. | Pérez-Martín et al. (2014) |
| ***5´CBLP*** | 5´-CTTCTCGCCCATGACCAC-3´ | Forward primer for *CBLP* gene expression analysis by RT-qPCR. | Pérez-Martín et al. (2014) |
| ***3´CBLP*** | 5´-CCCACCAGGTTGTTCTTCAG-3´ | Reverse primer for *CBLP* gene expression analysis by RT-qPCR. | Pérez-Martín et al. (2014) |

Pérez-Martín M, Pérez-Pérez ME, Lemaire SD, Crespo JL (2014). Plant Physiol, 166 (2):977-1008.

**Table S3.** Statistical analysis from main figures.

| Figure | Experimental condition | Statistical analysis | n | *P* value |
|---|---|---|---|---|
| 1F | Oxygen production WT 0h vs WT 8h +GLA | One-way Anova | 4 | *P*<0.001 *** |
| 1F | Y(II) WT 0h vs WT 8h +GLA | One-way Anova | 4 | *P*<0.001 *** |
| 1F | $F_v/F_m$ WT 0h vs WT 8h +GLA | One-way Anova | 4 | *P*<0.001 *** |
| 1G | ROS levels WT 0h vs WT 0.5h +GLA | One-way Anova | 3 | *P*<0.001 *** |
| 1G | ROS levels WT 0h vs WT 2h +GLA | One-way Anova | 3 | *P*<0.001 *** |
| 1G | ROS levels WT 0h vs WT 4h +GLA | One-way Anova | 3 | *P*<0.001 *** |
| 1G | ROS levels WT 0h vs WT 8h +GLA | One-way Anova | 3 | *P*<0.001 *** |
| 1H | ROS levels WT cytosol 0h vs WT cytosol 0.5h +GLA | One-way Anova | 3 | *P*<0.001 *** |
| 1H | ROS levels WT cytosol 0h vs WT cytosol 4h +GLA | One-way Anova | 3 | *P*<0.001 *** |
| 1H | ROS levels WT cytosol 0h vs WT cytosol 8h +GLA | One-way Anova | 3 | *P*<0.001 *** |
| 1H | ROS levels WT stroma 0h vs WT stroma 0.5h +GLA | One-way Anova | 3 | *P*<0.001 *** |
| 1H | ROS levels WT stroma 0h vs WT stroma 4h +GLA | One-way Anova | 3 | *P*<0.001 *** |
| 1H | ROS levels WT stroma 0h vs WT stroma 8h +GLA | One-way Anova | 3 | *P*<0.01 ** |
| 3A | ROS levels WT 0h dark vs *prk* 0h dark | Two-tailed Student´s *t* test | 4 | *P*≥0.05 Not significant |
| 3A | ROS levels WT 48h dark vs *prk* 48h dark | Two-tailed Student´s *t* test | 4 | *P*<0.05 * |
| 3A | ROS levels WT 0h LL vs *prk* 0h LL | Two-tailed Student´s *t* test | 4 | *P*<0.01 ** |
| 3A | ROS levels WT 48h LL vs *prk* 48h LL | Two-tailed Student´s *t* test | 4 | *P*<0.001 *** |
| 3A | ROS levels WT 0h ML vs *prk* 0h ML | Two-tailed Student´s *t* test | 4 | *P*<0.001 *** |
| 3A | ROS levels WT 48h ML vs *prk* 48h ML | Two-tailed Student´s *t* test | 4 | *P*<0.01 ** |
| 3C | ROS levels *prk* control 48h vs *prk* 56h control | Two-tailed Student´s *t* test | 3 | *P*<0.05 ** |
| 3C | ROS levels *prk* control 48h vs *prk* 56h +NAC | Two-tailed Student´s *t* test | 3 | *P*<0.05 ** |
| 3C | ROS levels *prk* control 48h vs *prk* 56h +acetate | Two-tailed Student´s *t* test | 3 | *P*<0.05 * |
| 3D | Oxygen production WT 52h control vs WT 52h +acetate | Two-tailed Student´s *t* test | 3 | *P*<0.01 ** |
| 3D | Oxygen production *prk* 52h control vs *prk* 52h +acetate | Two-tailed Student´s *t* test | 3 | *P*<0.01 ** |
| 3D | Y(II) WT 52h control vs WT 52h +acetate | Two-tailed Student´s *t* test | 3 | *P*≥0.05 Not significant |
| 3D | Y(II) *prk* 52h control vs *prk* 52h +acetate | Two-tailed Student´s *t* test | 3 | *P*<0.001 *** |
| 3E | Oxygen consumption WT 52h control vs WT 52h +acetate | Two-tailed Student´s *t* test | 3 | *P*<0.01 ** |
| 3E | Oxygen consumption WT 52h control vs WT 52h +acetate | Two-tailed Student´s *t* test | 3 | *P*<0.05 * |
| 4B | mRNA levels WT vs *atg8* | Two-tailed Student´s *t* test | 3 | *P*<0.001 *** |

| | | | | |
|---|---|---|---|---|
| 4D | Protein levels RPS6 WT vs *atg8* | Two-tailed Student´s *t* test | 3 | *P*<0.05 * |
| 4D | Protein levels RPL37 WT vs *atg8* | Two-tailed Student´s *t* test | 3 | *P*<0.001 *** |
| 4F | Chlorophyll content WT exp vs *atg8* exp | Two-tailed Student´s *t* test | 3 | *P*<0.05 * |
| 4F | Chlorophyll content WT stat vs *atg8* stat | Two-tailed Student´s *t* test | 3 | *P*<0.05 *** |
| 4G | ROS levels WT exp vs *atg8 exp* | Two-tailed Student´s *t* test | 3 | *P*<0.001 *** |
| 4G | ROS levels WT stat vs *atg8* stat | Two-tailed Student´s *t* test | 3 | *P*<0.05 * |
| 5B | Dead cells percentage WT 72h +GLA vs *atg8* 72h +GLA | Two-tailed Student´s *t* test | 4 | *P*<0.001 *** |
| 5C | ROS levels WT 0h vs *atg8* 0h | Two-tailed Student´s *t* test | 3 | *P*<0.001 *** |
| 5C | ROS levels WT 8h +GLA vs *atg8* 8h +GLA | Two-tailed Student´s *t* test | 3 | *P*<0.05 * |
| 5D | Oxygen production WT 24h +GLA vs *atg8* 24h +GLA | Two-tailed Student´s *t* test | 3 | *P*<0.05 * |
| 5D | Oxygen production WT 48h +GLA vs *atg8* 48h +GLA | Two-tailed Student´s *t* test | 3 | *P*<0.001 ** |
| 5D | Y(II) WT 24h +GLA vs *atg8* 24h +GLA | Two-tailed Student´s *t* test | 3 | *P*<0.001 *** |
| 5D | Y(II) WT 48h +GLA vs *atg8 48*h +GLA | Two-tailed Student´s *t* test | 3 | *P*<0.001 *** |
| 5D | $F_v/F_m$ WT 48h +GLA vs *atg8* 48h +GLA | Two-tailed Student´s *t* test | 3 | *P*<0.001 ** |
| 5D | Oxygen consumption WT 48h +GLA vs *atg8* 48h +GLA | Two-tailed Student´s *t* test | 3 | *P*<0.001 ** |
| 6A | $OD_{750nm}$ WT LL vs *atg8* LL | Two-tailed Student´s *t* test | 4 | *P*≥0.05 Not significant |
| 6A | $OD_{750nm}$ WT LL vs *prk* LL | Two-tailed Student´s *t* test | 4 | *P*≥0.05 Not significant |
| 6A | $OD_{750nm}$ WT LL vs *atg8 prk #2* LL | Two-tailed Student´s *t* test | 4 | *P*≥0.05 Not significant |
| 6A | $OD_{750nm}$ WT LL vs *atg8 prk #3* LL | Two-tailed Student´s *t* test | 4 | *P*≥0.05 Not significant |
| 6A | $OD_{750nm}$ WT ML vs *atg8* ML | Two-tailed Student´s *t* test | 4 | *P*≥0.05 Not significant |
| 6A | $OD_{750nm}$ WT ML vs *prk* ML | Two-tailed Student´s *t* test | 4 | *P*<0.05 ** |
| 6A | $OD_{750nm}$ WT ML vs *atg8 prk #2* ML | Two-tailed Student´s *t* test | 4 | *P*<0.05 ** |
| 6A | $OD_{750nm}$ WT ML vs *atg8 prk #3* ML | Two-tailed Student´s *t* test | 4 | *P*<0.001 ** |
| 6A | $OD_{750nm}$ WT HL vs *atg8* HL | Two-tailed Student´s *t* test | 4 | *P*<0.001 ** |
| 6A | $OD_{750nm}$ WT HL vs *prk* HL | Two-tailed Student´s *t* test | 4 | *P*<0.001 *** |
| 6A | $OD_{750nm}$ WT HL vs *atg8 prk #2* HL | Two-tailed Student´s *t* test | 4 | *P*<0.001 *** |
| 6A | $OD_{750nm}$ WT HL vs *atg8 prk #3* HL | Two-tailed Student´s *t* test | 4 | *P*<0.001 *** |

*** *P*<0.001; ** *P*<0.01; **P*<0.05; Not significant ≥0.05

**Table S4.** Statistical analysis from supplemental figures.

| Figure | Experimental condition | Statistical analysis | n | *P* value |
|---|---|---|---|---|
| Suppl. S1B | Ru5P/Xu5P WT 0h vs WT 0.5h +GLA | One-way Anova | 3 | *P*<0.01 ** |
| Suppl. S1B | Ru5P/Xu5P WT 0h vs WT 1h +GLA | One-way Anova | 3 | *P*<0.001 *** |
| Suppl. S1B | Ru5P/Xu5P WT 0h vs WT 2h +GLA | One-way Anova | 3 | *P*<0.001 *** |
| Suppl. S1B | Ru5P/Xu5P WT 0h vs WT 4h +GLA | One-way Anova | 3 | *P*<0.001 *** |
| Suppl. S1B | Ru5P/Xu5P WT 0h vs WT 8h +GLA | One-way Anova | 3 | *P*<0.001 *** |
| Suppl. S3A | NADPH fluorescence WT 0h vs WT 8h +GLA | One-way Anova | 3 | *P*<0.01 ** |
| Suppl. S3B | Oxygen consumption WT 0h vs WT 8h +GLA | One-way Anova | 3 | *P*<0.001 *** |
| Suppl. S5A | Ru5P/Xu5P WT 48h vs *prk* 48h | Two-tailed Student´s *t* test | 3 | *P*<0.01 ** |
| Suppl. S5A | RuBP WT 48h vs *prk* 48h | Two-tailed Student´s *t* test | 3 | *P*<0.01 ** |
| Suppl. S5A | 3PGA WT 48h vs *prk* 48h | Two-tailed Student´s *t* test | 3 | *P*<0.01 ** |
| Suppl. S5A | G3P WT 48h vs *prk* 48h | Two-tailed Student´s *t* test | 3 | *P*<0.05 * |
| Suppl. S5A | E4P WT 48h vs *prk* 48h | Two-tailed Student´s *t* test | 3 | *P*<0.05 * |
| Suppl. S5A | R5P WT 48h vs *prk* 48h | Two-tailed Student´s *t* test | 3 | *P*<0.01 ** |
| Suppl. S5B | NADPH WT 48h vs *prk* 48h | Two-tailed Student´s *t* test | 3 | *P*<0.01 ** |
| Suppl. S5C | DHAP WT 48h vs *prk* 48h | Two-tailed Student´s *t* test | 3 | *P*<0.001 *** |
| Suppl. S5C | PEP WT 48h vs *prk* 48h | Two-tailed Student´s *t* test | 3 | *P*<0.001 *** |
| Suppl. S5C | Pyr WT 48h vs *prk* 48h | Two-tailed Student´s *t* test | 3 | *P*<0.001 *** |
| Suppl. S5C | Acetyl-CoA WT 48h vs *prk* 48h | Two-tailed Student´s *t* test | 3 | *P*<0.01 ** |
| Suppl. S5D | Citrate WT 48h vs *prk* 48h | Two-tailed Student´s *t* test | 3 | *P*<0.001 *** |
| Suppl. S5D | Isocitrate WT 48h vs *prk* 48h | Two-tailed Student´s *t* test | 3 | *P*<0.01 ** |
| Suppl. S5D | Succinate WT 48h vs *prk* 48h | Two-tailed Student´s *t* test | 3 | *P*<0.001 *** |
| Suppl. S5D | Fumarate WT 48h vs *prk* 48h | Two-tailed Student´s *t* test | 3 | *P*<0.01 ** |
| Suppl. S5D | Malate WT 48h vs *prk* 48h | Two-tailed Student´s *t* test | 3 | *P*<0.01 ** |
| Suppl. S6A | Acetate WT 0h vs WT 24h | One-way Anova | 3 | *P*<0.001 *** |
| Suppl. S6A | Acetate WT 0h vs WT 48h | One-way Anova | 3 | *P*<0.001 *** |
| Suppl. S6A | Acetate *prk* 0h vs *prk* 24h | One-way Anova | 3 | *P*<0.001 *** |
| Suppl. S6A | Acetate *prk* 0h vs *prk* 48h | One-way Anova | 3 | *P*<0.001 *** |

| | | | | |
|---|---|---|---|---|
| Suppl. S7A | $F_v/F_m$ WT 0h vs *prk* 52h | One-way Anova | 3 | $P \geq 0.05$ Not significant |
| Suppl. S7A | $F_v/F_m$ *prk* 0h vs *prk* 52h | One-way Anova | 3 | $P \geq 0.05$ Not significant |

*** $P<0.001$; ** $P<0.01$; *$P<0.05$; Not significant ≥0.05
